# Gene duplication of SNAPC1 generates transcription factors for snRNAs and sex-specific piRNAs

**DOI:** 10.64898/2026.08.31.747621

**Authors:** Lars Kristian Benner, Margaret R Starostik, Rebecca J Tay, Ayaka Inoki, Mindy Clark, Suhua Feng, Michael C Schatz, Steven E Jacobsen, John K Kim

## Abstract

Piwi-interacting RNAs (piRNAs) are small non-coding RNAs essential for transposon silencing and germline integrity across metazoans. In many species, piRNA expression is sexually dimorphic, yet the molecular mechanisms underlying this sex specificity remain poorly understood. In *Caenorhabditis elegans,* sexually dimorphic piRNA expression is regulated at the transcriptional level. We previously identified SNPC-1.3, a paralog of the small nuclear RNA (snRNA) activating protein complex (SNAPc/SNPC) subunit SNAPC1, as a male-specific piRNA transcription factor. However, the factors governing female piRNA expression remained elusive. Here, we identify SNPC-1.2, a second SNPC-1 paralog, as a female-specific piRNA transcription factor. SNPC-1.2 interacts with the core piRNA transcriptional machinery, binds female piRNA loci, is required for female piRNA expression, and promotes hermaphrodite fertility. In contrast, a third paralog, SNPC-1.1, retains the ancestral SNAPc function in snRNA transcription and is dispensable for piRNA biogenesis. Together, these findings reveal how gene duplication and functional specialization within the *snpc-1* gene family generate specificity factors that direct the core SNAP complex to distinct genomic targets, providing a molecular mechanism for sexually dimorphic piRNA expression while maintaining canonical snRNA transcription.

## Introduction

Piwi-interacting RNAs (piRNAs) comprise a class of germline-enriched small RNAs that silence foreign DNA elements such as transposable elements, which can disrupt gene coding regions and induce DNA breaks (Gasior et al., 2006; Vagin et al., 2006; Aravin et al., 2007; Brennecke et al., 2007). Small RNAs function by associating with Argonaute proteins (reviewed in Luteijn and Ketting, 2013; Iwasaki et al., 2015; Ozata et al., 2019), with piRNAs specifically associating with the PIWI subfamily of Argonautes. Loss of piRNAs leads to severe germline defects in a variety of species, underscoring their importance for ensuring animal fertility (Lin and Spradling, 1997; Cox et al., 1998; Deng and Lin, 2002; Girard et al., 2006; Houwing et al., 2007; Wang and Reinke, 2008; Kumar et al., 2019).

Sex-specific or sex-biased piRNA expression has been reported across metazoans (Zhou et al., 2010; Billi et al., 2013; Yang et al., 2013; Williams et al., 2015), and is often linked to biological function. For example, in the silkworm *Bombyx mori*, a female-specific piRNA controls sex determination, whereas in *Drosophila,* sex-biased piRNA populations mirror sex-specific transposon activity (Kiuchi et al., 2014; Chen et al., 2021).

The nematode *Caenorhabditis elegans* provides a powerful system to dissect sex-specific piRNA regulation. As hermaphrodites, these animals undergo spermatogenesis during the final L4 larval stage, then switch to oogenesis in adulthood, enabling the study of sex-specific piRNA expression within a single germline. Distinct piRNA subsets are enriched during spermatogenesis (“male piRNAs”) or oogenesis (“female piRNAs”), while others show no significant enrichment between stages (“not significantly enriched” or NE piRNAs). Because male and female piRNA genes are associated with distinct upstream *cis*-regulatory elements, their differential expression is likely controlled at the transcriptional level (Billi et al., 2013). However, the molecular mechanisms that drive sex-specific piRNA transcription remain poorly understood.

Recent studies in *C. elegans* have begun to illuminate the mechanisms regulating piRNA transcription, providing a framework for understanding how sex-specific expression may be achieved. In *C. elegans*, piRNAs are 21 nucleotides (nt) long with a characteristic uracil bias at the first position (i.e. 21U RNAs). Most of these 21U piRNAs are transcribed from two large megabase-sized clusters on chromosome IV (Ruby et al., 2006). Each piRNA is transcribed as an independent unit marked by an upstream 8-nt core motif, termed the “Ruby motif,” located approximately 40 base pairs (bp) upstream of a YRNT motif that defines the transcription start site (Ruby et al., 2006).

Several trans-acting factors have been identified that recognize these cis-regulatory elements. The core piRNA transcription factor SNPC-4 (Kasper et al., 2014), together with TOFU-4, TOFU-5 (Weng et al., 2019), and the putative kinase PRDE-1 (Weick et al., 2014; Kasper et al., 2014; Weng et al., 2019), forms an upstream sequence transcription complex (USTC) that promotes piRNA expression (Weng et al., 2019). However, how the composition or regulation of this machinery differs at male- and female-specific piRNA loci remains largely unknown.

*C. elegans* SNPC-4 is a core component of the small nuclear (snRNA) activating protein complex (SNAPc), a highly conserved complex found from trypanosomes to humans. The minimal SNAPc in humans comprises three subunits, SNAPC4, SNAPC3, and SNAPC1, analogous to SNAP190, SNAP50, SNAP43 in *Drosophila*, respectively (Mittal et al., 1999; Ma and Hernandez, 2002; Hinkley et al., 2003; Jawdekar et al., 2006). These subunits assemble in a 1:1:1 stoichiometry before binding the proximal sequence element (PSE) and recruiting RNA polymerase II or III for snRNA transcription (Lai et al., 2008; Hung and Stumph, 2012).

Among these subunits, SNAPC1 remains the least characterized. Although it lacks similarity to other known proteins, its N-terminal SNAPC1 domain is highly conserved (Hung and Stumph, 2011), and structure-function analyses have been conducted in humans and *Drosophila* (Ma and Hernandez, 2001; Hung et al., 2009; Sun et al., 2022). *C. elegans* presents a unique opportunity to study SNAPC1 functional diversification. Unlike flies and humans, which each encode a single *SNAPC1* gene, *C. elegans* encodes four SNAPC1 paralogs: SNPC-1.1, SNPC-1.2, SNPC-1.3, and SNPC-1.4 (Fig. 1A). Whether these paralogs have diverged functionally remains an open question.

**Figure 1.**
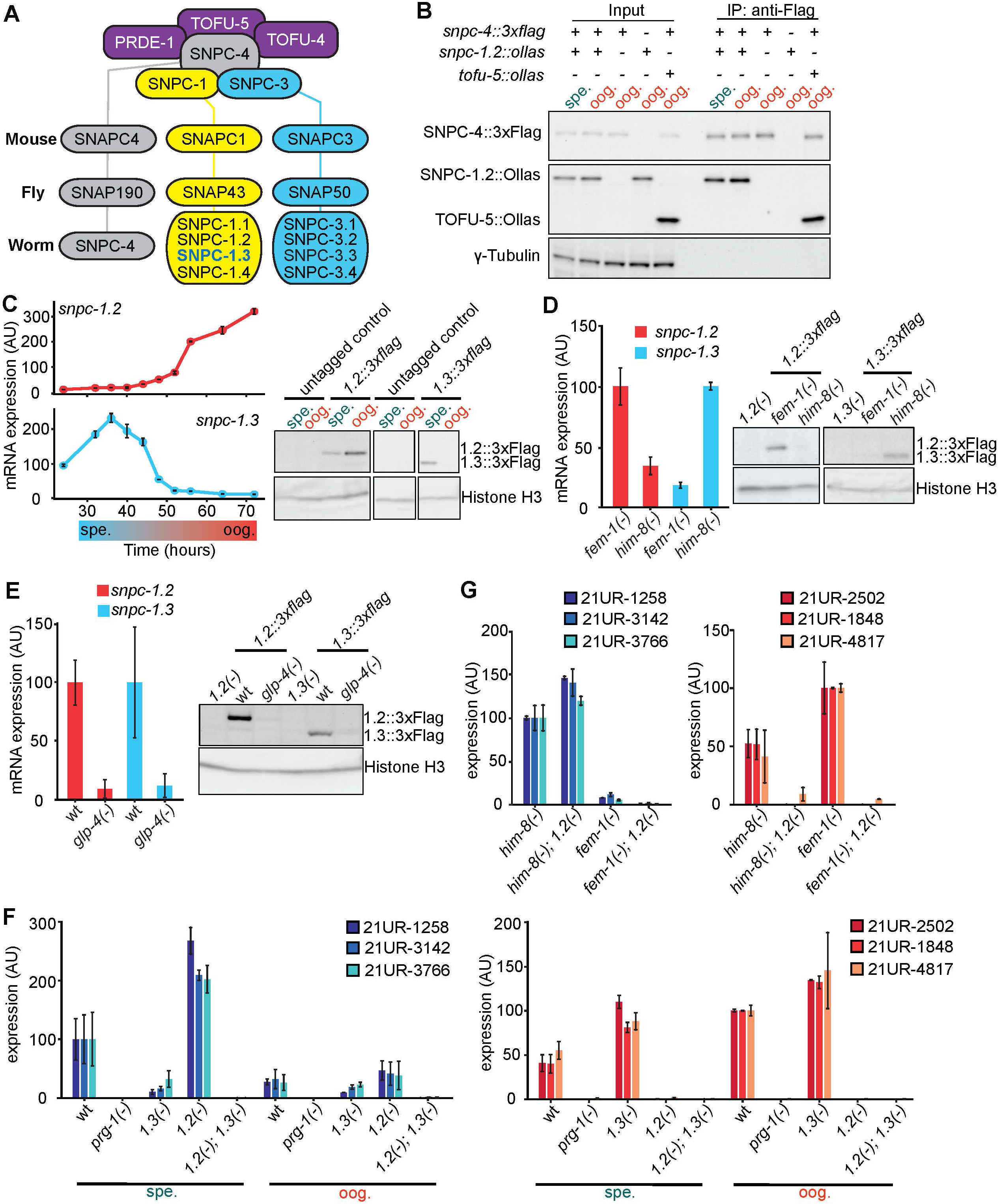
SNPC-1.2 is expressed in the female germline and is required for female piRNA transcription. **(A)** Schematic of the conserved heterotrimeric SNAPc (SNPC) complex consisting of SNPC-1 (yellow), SNPC-3 (blue), and SNPC-4 (gray). The previously identified male-specific piRNA transcription factor SNPC-1.3 is highlighted (blue text; Choi et al., 2021). Known piRNA factors comprising the Upstream Transcription Complex (USTC) that interact with the *C. elegans* SNPC complex—TOFU-4, TOFU-5, and PRDE-1—are shown in purple (Weick et al., 2014; Weng et al., 2019). Homologs of SNPC subunits from *H. sapiens*, *D. melanogaster*, and *C. elegans* are indicated. **(B)** SNPC-4 interacts with SNPC-1.2. Anti-Flag immunoprecipitation of SNPC-4::3xFlag during spermatogenesis (spe.) and oogenesis (oog.). TOFU-5::Ollas serves as a positive control for a SNPC-4::3xFlag interactor (Weng et al., 2019). γ-tubulin is shown as a loading control. **(C)** *snpc-1.2* mRNA and protein are enriched during oogenesis. Left: qPCR time course of *snpc-1.2* and *snpc-1.3* mRNA expression across development, normalized to *eft-2*. Error bars represent ± SD of two technical replicates. Right: Western blot of SNPC-1.2::3xFlag and SNPC-1.3::3xFlag during spermatogenesis and oogenesis. Histone H3 serves as a loading control. **(D)** *snpc-1.2* mRNA and protein are enriched in females. Left: qPCR of *snpc-1.2* and *snpc-1.3* mRNA in *fem-1(-)* females and *him-8(-)* males, normalized to *eft-2*. Error bars represent ± SD of two technical replicates. Right: Western blot of SNPC-1.2::3xFlag and SNPC-1.3::3xFlag in *fem-1(-)* females and *him-8(-)* males. **(E)** *snpc-1.2* mRNA and protein are germline enriched. Left: qPCR of *snpc-1.2* and *snpc-1.3* mRNA in wild-type and germline-deficient *glp-4(-)* animals at 25°C. Error bars represent ± SD of two technical replicates. Right: Western blot of SNPC-1.2::3xFlag and SNPC-1.3::3xFlag in wild type and *glp-4(-)* mutants at 25°C. **(F)** SNPC-1.2 is required for female piRNA expression. TaqMan qPCR of male (left) and female (right) piRNAs at spermatogenesis (spe.) and oogenesis (oog.). Expression normalized to U18 small nucleolar RNA (snoRNA). Error bars represent ± SD of two technical replicates. **(G)** SNPC-1.2 is required for female piRNA expression in both sexes. TaqMan qPCR of male and female piRNAs normalized to U18 snoRNA. Error bars represent ± SD of two technical replicates.

We previously identified SNPC-1.3 as a male-specific piRNA transcription factor (Choi et al., 2021). Here, we identify SNPC-1.2 as its female counterpart and SNPC-1.1 as a distinct SNPC-1 paralog that promotes snRNA transcription. These findings demonstrate that robust and specific expression of sex-specific piRNAs and snRNAs in *C. elegans* is mediated through the duplication and functional divergence of the ancient SNAPC1/SNPC-1 transcription factor family.

## Results

### SNPC-1.2 interacts with the shared piRNA and snRNA transcription factor SNPC-4

Having previously identified SNPC-1.3 as a male-specific piRNA transcription factor, we hypothesized a female-specific counterpart may exist. To explore this possibility, we reanalyzed our SNPC-4 immunoprecipitation and mass spectrometry datasets from masculinized *him-8(-)* and feminized *fem-1(-)* worms (Choi et al., 2021; Hodgkin et al., 1979; Doniach and Hodgkin, 1984), focusing on proteins enriched in the feminized background. Peptides corresponding to SNPC-1.2 were significantly enriched in *fem-1(-)* relative to *him-8(-)* samples, suggesting a potential female-specific role for SNPC-1.2 in piRNA regulation.

To validate the interaction between SNPC-1.2 and SNPC-4 identified by mass spectrometry, we used CRISPR/Cas9 genome editing (Paix et al., 2015) to tag the C-terminus of SNPC-1.2 with an Ollas epitope. We crossed the *snpc-1.2::ollas* allele into a *snpc-4::3xflag* background and performed anti-Flag immunoprecipitations followed by anti-Ollas immunoblotting at 48 and 72 hours post-L1 hatching at 20°C, corresponding to spermatogenesis and oogenesis, respectively. SNPC-4::3xFlag co-immunoprecipitated with SNPC-1.2::Ollas at both time points (Fig. 1B).

To further confirm the interaction, we swapped the Flag and Ollas epitopes on SNPC-1.2 and SNPC-4 by CRISPR-Cas9, generating a strain carrying a C-terminal 3xFlag tag on SNPC-1.2 and an Ollas tag on SNPC-4. The resulting *snpc-1.2::3xflag* and *snpc-4::ollas* alleles were crossed to create a strain expressing both tagged proteins from their endogenous loci. Reciprocal co-immunoprecipitation again revealed robust interaction at both 48 and 72 hour time points, consistent with SNPC-1.2 and SNPC-4 associating during both spermatogenesis and oogenesis (Supplemental Fig. S1A). The 3xFlag tag on SNPC-1.2 did not impair piRNA expression, confirming that the tagged protein retains wild-type function (Supplemental Fig. S1B).

### SNPC-1.2 is upregulated in the female germline

Although SNPC-1.2 interacts with SNPC-4 during both spermatogenesis and oogenesis, we asked whether *snpc-1.2* expression differs between these stages. Using mRNA qPCR, we found that *snpc-1.2* transcripts are highly upregulated during oogenesis (Fig. 1C). As a control, we examined *snpc-1.3* mRNA expression, which is almost exclusively expressed during spermatogenesis but not oogenesis (Choi et al., 2021). Protein expression analysis similarly showed that SNPC-1.2::3xFlag is elevated during oogenesis, whereas SNPC-1.3::3xFlag is robustly detected only during spermatogenesis (Fig. 1C). These results indicate that SNPC-1.2 is expressed at both stages but significantly upregulated during oogenesis.

To further examine the enrichment of SNPC-1.2 expression in the female germline, we compared its mRNA and protein levels in masculinized *him-8(-)* males and feminized *fem-1(-)* females. While *snpc-1.3* mRNA was highly enriched in *him-8(-)* males, *snpc-1.2* mRNA was most strongly expressed in *fem-1(-)* females (Fig. 1D). Consistent with transcript levels, SNPC-1.2::3xFlag protein was enriched in females, while SNPC-1.3::3xFlag was enriched in males (Fig. 1D), supporting the idea that SNPC-1.2 functions preferentially during oogenesis.

Most piRNA pathway proteins are germline-enriched (Cox et al., 1998; Reinke et al., 2000; Batista et al., 2008; de Albuquerque et al., 2014; Kasper et al., 2014; Weick et al., 2014; Choi et al., 2021). We therefore tested whether SNPC-1.2 expression is also germline-dependent. We measured *snpc-1.2* and *snpc-1.3* mRNA levels in wildtype and *glp-4(-)* mutants, which lack a mature germline at 25°C (Beanan and Strome, 1992). At this restrictive temperature, both transcripts were strongly reduced in *glp-4(-)* animals compared to wild-type. Correspondingly, SNPC-1.2::3xFlag protein, like SNPC-1.3::3xFlag, was nearly undetectable in *glp-4(-)* mutants (Fig. 1E). These findings demonstrate that SNPC-1.2 is a germline-enriched transcription factor preferentially expressed in the female germline.

### SNPC-1.2 is required for female piRNA transcription

Given that SNPC-1.2 is germline-expressed and interacts with the piRNA transcription factor SNPC-4, we next tested whether it is required for piRNA biogenesis. We used CRISPR-Cas9 genome editing to generate a 112-bp deletion at the *snpc-1.2* locus that introduces a premature stop codon within the N-terminal region (Supplemental Fig. S1C). To assess piRNA expression, we quantified several representative male and female piRNAs (Billi et al., 2013) in the *snpc-1.2(-)* mutant by TaqMan qPCR, alongside the piRNA-loading Piwi mutant *prg-1(-)* and the male piRNA transcription factor mutant *snpc-1.3(-)* as controls. Loss of *snpc-1.2* nearly eliminated female piRNA expression during both spermatogenesis (when they are normally low) and oogenesis (when they are highly expressed). In contrast, male piRNAs were modestly elevated during spermatogenesis in the *snpc-1.2(-)* mutant (Fig. 1F).

To further examine the role of *snpc-1.2* on female piRNA expression, we generated a second *snpc-1.2(-)* mutant allele using two independent crRNAs. This alternative allele (Supplemental Fig. S1C) introduces a premature stop codon and similarly results in a selective loss of female piRNAs, further supporting a requirement for SNPC-1.2 in female piRNA biogenesis (Supplemental Fig. S1D). To test whether this effect was consistent across germline sexual states, we measured piRNA expression in *snpc-1.2(-)* mutants in both *him-8(-)* males and *fem-1(-)* females. In both contexts, female piRNAs were dramatically depleted, whereas *him-8(-); snpc-1.2(-)* males showed a slight upregulation of male piRNAs (Fig. 1G). Finally, in the *snpc-1.2(-); snpc-1.3(-)* double mutant, both male and female piRNAs were nearly undetectable at all stages (Fig. 1F), indicating that SNPC-1.2 and SNPC-1.3 together are required for sex-specific piRNA expression.

To assess the global impact of SNPC-1.2 on piRNA expression, we performed small RNA sequencing in *snpc-1.2(-)* mutants. We generated additional wild-type, *snpc-1.2(-)*, *snpc-1.3(-)*, and *snpc-1.2(-); snpc-1.3(-)* small RNA datasets collected during spermatogenesis and oogenesis, substantially expanding our previous analysis of piRNAs (Choi et al., 2021). We also developed a customized piRNA quantification pipeline based on HTSeq (Anders et al., 2015), optimized to distinguish individual piRNAs within dense clusters and identify novel piRNAs that may have been previously misannotated (see Methods). Using this pipeline (Supplemental Fig. S2A), we identified 1,763 novel piRNAs: 48% (851) mapped to chromosome IV (Fig. 2A), and 16% overlapped with recently reported novel piRNAs (Seroussi et al., 2023) (Fig. 2B).

**Figure 2.**
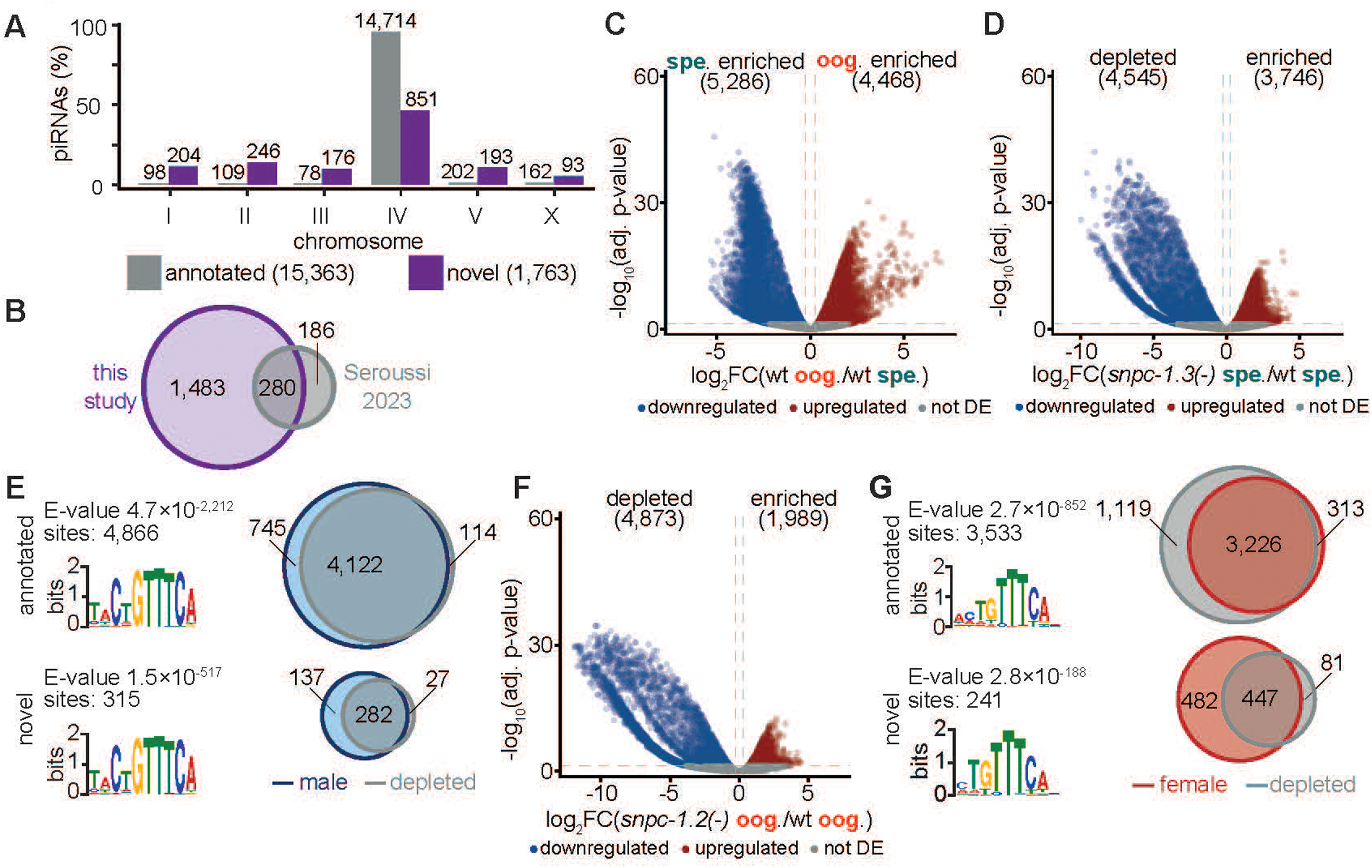
SNPC-1.2 is required for transcription of female piRNAs. **(A)** Chromosomal distribution of annotated and newly identified piRNAs. Numbers above bars indicate total piRNAs per chromosome. **(B)** Overlap of newly identified piRNAs with those reported by Seroussi et al. (2023). **(C)** Differential expression of piRNAs during spermatogenesis (spe.) and oogenesis (oog.) in wild-type animals. **(D)** Loss of *snpc-1.3* reduces male piRNA expression. Volcano plot of differential expression of piRNAs during spermatogenesis in *snpc-1.3(-)* mutants compared with wild type. **(E)** Male piRNAs are depleted in *snpc-1.3(-)* mutants. Top: Annotated piRNAs. Bottom: Novel piRNAs. Left: Sequence logos of the conserved upstream motifs for male piRNAs. Right: Venn diagrams showing overlap of spermatogenesis-enriched piRNAs in wild type and piRNAs depleted in *snpc-1.3(-)* mutants during spermatogenesis. **(F)** SNPC-1.2 is required for female piRNA expression. Volcano plot showing differential expression of piRNAs during oogenesis in *snpc-1.2(-)* mutants compared with wild type. **(G)** Female piRNAs are depleted in *snpc-1.2(-)* mutants. Top: Annotated piRNAs. Bottom: Novel piRNAs Left: Sequence logos of the conserved upstream motifs for female piRNAs. Right: Venn diagrams showing overlap between oogenesis-enriched piRNAs in wild type and piRNAs depleted in *snpc-1.2(-)* mutants during oogenesis.

With the expanded dataset and updated annotation, we identified two major piRNA populations in wild-type hermaphrodites: 5,286 enriched during spermatogenesis (“male piRNAs”) and 4,468 enriched during oogenesis (“female piRNAs”) (Fig. 2C; Supplemental Table 1). Consistent with previous findings (Choi et al., 2021), male piRNAs were significantly depleted in *snpc-1.3(-)* mutants relative to wild type (Supplemental Table 2, 3). Of 5,286 male piRNAs, 83% were significantly downregulated in *snpc-1.3(-)* mutants (Fig. 2D; Supplemental Table 2), including 4,122 annotated and 282 novel male piRNAs (Fig. 2E). These novel male piRNAs also exhibited a strong 5’ cytosine bias at the first nucleotide position of their upstream Ruby motifs (Fig. 2E), consistent with prior studies (Billi et al., 2013; Choi et al., 2021; Ruby et al., 2006).

We next examined how loss of *snpc-1.2* affects female piRNA expression (Supplemental Table 4, 5). Of the 4,468 piRNAs enriched during oogenesis in wild-type hermaphrodites, 82% were significantly downregulated in *snpc-1.2(-)* mutants, including 3,226 previously annotated and 447 novel female piRNAs (Fig. 2F, G; Supplemental Table 5). Analysis of the upstream sequences confirmed the presence of the Ruby motif but revealed no strong 5’ cytosine bias, in agreement with earlier observations for female piRNAs (Fig. 2G) (Billi et al., 2013; Choi et al., 2021).

Finally, to evaluate the combined effects of *snpc-1.2* and *snpc-1.3* loss, we analyzed small RNA-seq data from *snpc-1.2(-); snpc-1.3(-)* double mutants (Supplemental Table 6, 7). During spermatogenesis, 3,965 piRNAs were depleted, including 62% male and 32% female piRNAs (Supplemental Table 6). During oogenesis, 2,873 piRNAs were downregulated, 75% of which were female and 16% male, relative to wild type (Supplemental Fig. S2B; Supplemental Table 7). The preferential loss of both male and female piRNAs during spermatogenesis, but primarily female piRNAs during oogenesis, in *snpc-1.2(-); snpc-1.3(-)* double mutants is consistent with SNPC-1.3 being expressed almost exclusively during spermatogenesis, whereas SNPC-1.2 is more broadly expressed with peak levels during oogenesis (Fig. 1C). Together, our expanded small RNA-seq datasets and improved annotation indicate that SNPC-1.2 globally promotes female piRNA expression and partially supports male piRNA transcription in the absence of SNPC-1.3.

### SNPC-1.4 modestly promotes male and female piRNA expression

Since SNPC-1.4 interacts with SNPC-4 (Choi et al., 2021), we wanted to explicitly test the role of *snpc-1.4* in piRNA biogenesis. We used CRISPR-Cas9 genome editing to generate a *snpc-1.4(-)* null allele by inserting a non-endogenous sequence resulting in a premature termination codon in the second exon (Supplemental Fig. S1C). We then performed TaqMan qPCR measuring several male and female piRNA levels in the *snpc-1.4(-)* mutant. The *snpc-1.4(-)* mutant worms showed only a small decrease in both male and female piRNA levels compared to wildtype at spermatogenesis and oogenesis (Supplemental Fig. S1E).

Since SNPC-4 is known to transcribe other classes of small RNAs including snRNAs, we tested if SNPC-1.4 is responsible for the transcription of these small RNA classes. We conducted qPCR of the *U1* snRNA in *snpc-1.4(-)* mutants and again observed only a modest decrease at both spermatogenesis and oogenesis compared to wildtype (Supplemental Fig. S1F). Thus, SNPC-1.4 does not significantly affect snRNA expression.

### SNPC-1.2 preferentially binds female piRNA loci in a SNPC-4-dependent manner

Given the role of SNPC-1.2 in promoting female piRNA expression, we hypothesized that SNPC-1.2 preferentially binds female piRNA promoters. We first tested whether SNPC-1.2 associates with piRNA clusters using anti-Flag ChIP-qPCR in the *snpc-1.2::3xflag* and wild-type (no-tag control) strains during oogenesis. We also examined whether SNPC-4 is required for SNPC-1.2 binding by depleting SNPC-4 via the auxin-inducible degron (AID) system. We added an AID tag to the C-terminus of SNPC-4 using CRISPR/Cas9 and crossed this strain into worms expressing TIR1 under the germline promoter *sun-1* (*snpc-1.2::3xflag; snpc-4::aid::ollas; Psun-1::TIR1*). TIR1 mediates auxin-dependent degradation of AID-tagged proteins, thereby depleting germline SNPC-4::AID upon auxin addition. As expected, SNPC-1.2 bound robustly to piRNA clusters on chromosome IV, and this binding was lost upon SNPC-4 depletion, while binding at an intergenic control region remained negligible (Fig. 3A).

**Figure 3.**
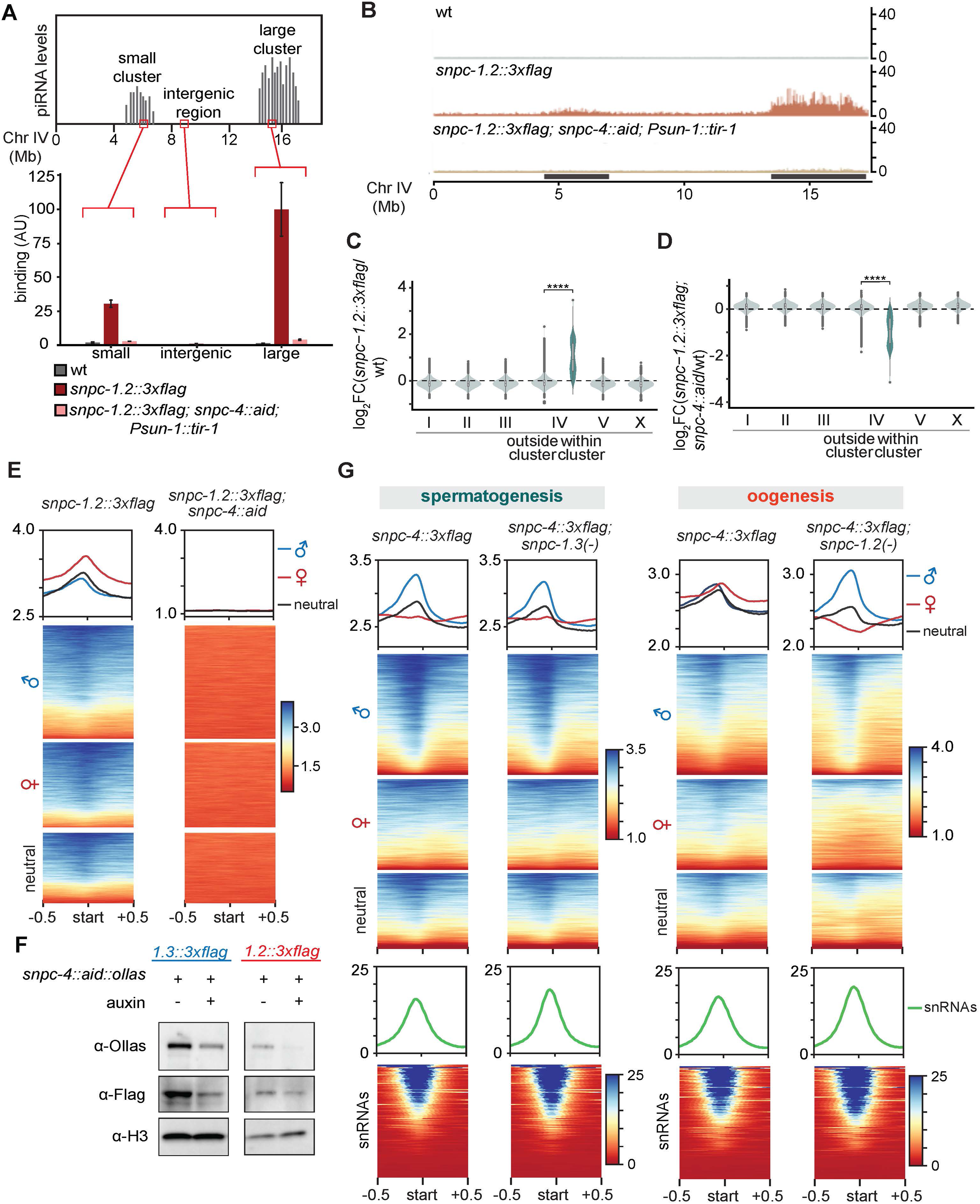
Reciprocal but asymmetric dependencies between SNPC-1 paralogs and SNPC-4 for piRNA loci binding. **(A)** SNPC-1.2 binds piRNA clusters in a SNPC-4-dependent manner. SNPC-1.2 binding on chromosome IV was assessed by ChIP-qPCR in wild type (N2), *snpc-1.2::3xflag*, and *snpc-1.2::3xflag; snpc-4::aid::ollas; Psun-1::TIR1* animals, in which SNPC-4::AID::Ollas is degraded upon auxin addition (250µM, 4 h). ChIP signal is normalized to input DNA (± SD of two technical replicates). Top: schematic of piRNAs transcribed from the small and large piRNA clusters on *C. elegans* chromosome IV. **(B)** SNPC-1.2 binding profiles across chromosome IV in N2, *snpc-1.2::3xflag*, and *snpc-1.2::3xflag; snpc-4::aid* animals. Shaded regions indicate the two piRNA clusters. **(C)** SNPC-1.2–bound regions are enriched within piRNA clusters relative to regions outside the piRNA clusters on chromosome IV (****p≤0.0001, Wilcoxon rank sum test). **(D)** Upon SNPC-4 depletion, SNPC-1.2 binding within piRNAs clusters is reduced relative to regions outside the piRNA clusters on chromosome IV (****p≤0.0001, Wilcoxon rank sum test). **(E)** SNPC-1.2 is enriched at piRNA transcription start sites. Metaplot shows the distribution of SNPC-1.2 reads (mean density ± standard error) centered on the 5′ nucleotide of mature piRNAs. Heat maps show ChIP signal around the 5′ nucleotide of mature piRNAs. **(F)** SNPC-4 depletion reduces SNPC-1.2 and SNPC-1.3 stability. Western blot of SNPC-1.3::3xFlag (left; spermatogenesis) and SNPC-1.2::3xFlag (right; oogenesis) following SNPC-4::AID::Ollas depletion with auxin (250 µM, 4 h). Histone H3 serves as a loading control. **(G)** SNPC-4 binds piRNA and snRNA loci genome-wide. During spermatogenesis, SNPC-4 preferentially binds male piRNA loci independently of *snpc-1.3*. During oogenesis, SNPC-4 binds both male and female piRNA loci, and binding at female piRNA loci requires *snpc-1.2*. Metaplot and heat map show enrichment of SNPC-4 ChIP-seq signal over male (blue), female (red), and NE (black) piRNA loci; snRNA loci (green) are shown below.

To define SNPC-1.2 binding genome-wide and assess its dependency on SNPC-4, we performed ChIP-seq of *snpc-1.2::3xflag*, *snpc-1.2::3xflag; snpc-4::aid; Psun-1::TIR1*, and wild-type (no-tag control) during oogenesis. SNPC-1.2 showed strong binding enrichment at the piRNA clusters on chromosome IV, with binding drastically reduced upon SNPC-4 depletion, confirming that SNPC-1.2 requires SNPC-4 for DNA association at piRNA loci (Fig. 3B). We quantified SNPC-1.2::3xFlag ChIP-seq signal over consecutive, non-overlapping 1 kb genomic bins across the genome. SNPC-1.2 binding was significantly enriched within the two piRNA clusters compared to regions outside the piRNA clusters on chromosome IV (Fig. 3C, D).

To determine whether SNPC-1.2 binds piRNA loci in a sex-specific manner, we compared ChIP-seq profiles at piRNA loci classified as male, female, or not enriched (NE) based on our small RNA-seq analysis (Fig. 2C). SNPC-1.3 binds almost exclusively to male piRNA loci in a SNPC-4-dependent manner during spermatogenesis (Supplemental Fig. S3A; Choi et al., 2021). SNPC-1.2 showed strongest binding at female piRNA loci, with progressively weaker binding at NE and male piRNA loci during oogenesis (Fig. 3E, Supplemental Fig. S3B). This binding was uniformly dependent on SNPC-4. Both SNPC-1.2 and SNPC-1.3 thus exhibit preferential binding to their respective piRNA classes, although SNPC-1.2 shows broader binding than SNPC-1.3.

To understand how SNPC-4 loss affects SNPC-1.2 DNA binding, we examined SNPC-1.2 and SNPC-1.3 protein levels with or without SNPC-4. Western blots revealed that depletion of SNPC-4 reduced, but did not eliminate, protein levels of both SNPC-1.2 and SNPC-1.3 (Fig. 3F), suggesting that SNPC-4 promotes the stability of these paralogs but also contributes to their DNA binding through mechanisms beyond simply maintaining protein abundance.

We next addressed the reciprocal question: how do SNPC-1.2 and SNPC-1.3 affect SNPC-4 binding to piRNA loci? To test this, we performed ChIP-seq on *snpc-4::3xflag*, *snpc-4::3xflag; snpc-1.3(-)*, and *snpc-4::3xflag; snpc-1.2(-)* worms. During spermatogenesis, loss of *snpc-1.3* did not affect SNPC-4 binding to male piRNA loci (Fig. 3G, Supplemental Fig. S3C, D), indicating that SNPC-4 can associate with these loci independently of SNPC-1.3. In contrast, loss of *snpc-1.2* substantially reduced SNPC-4 binding at female piRNA loci during oogenesis (Fig. 3G, Supplemental Fig. S3C, D), demonstrating that SNPC-1.2 is required for SNPC-4 recruitment to these sites. Importantly, SNPC-4 binding at snRNA loci was unaffected by the loss of either *snpc-1.2* or *snpc-1.3*, consistent with its independent role in snRNA biogenesis (Fig. 3G, Supplemental Fig. S3C, D). Together, these data show that SNPC-4 interacts with SNPC-1 paralogs in distinct, context-specific ways: it cooperates with SNPC-1.3 at male piRNA loci and with SNPC-1.2 at female piRNA loci, while functioning independently of these proteins in snRNA transcription. These findings suggest distinct regulatory modes: SNPC-4 is pre-bound at male piRNA loci and requires SNPC-1.3 for transcriptional activation, whereas SNPC-1.2 is necessary to recruit SNPC-4 to female piRNA promoters.

### Loss of *snpc-1.2* results in transgenerational subfertility without severe meiotic defects

To assess the physiological impact of female piRNA loss, we investigated fertility defects in *snpc-1.2(-)* mutants. In *prg-1(-)* mutants, destabilization of piRNAs results in temperature-sensitive fertility loss, particularly at 25°C (Wang and Reinke, 2008). Given that *snpc-1.2(-)* animals lose a large subset of piRNAs (Fig. 2F, Supplemental Fig. S7A, B), we hypothesized that they would also exhibit fertility defects. To test this, we shifted hermaphrodites to 25°C at the P0 stage and counted F1 progeny. Surprisingly, *snpc-1.2(-)* mutants did not show a marked reduction in fertility, whereas both *prg-1(-)* and *snpc-1.3(-)* mutants displayed significant fertility defects, consistent with previous reports (Supplemental Fig. S4A) (Batista et al., 2008; Wang and Reinke, 2008; Choi et al., 2021). Notably, the fertility defect of *snpc-1.3(-)* mutants was significantly enhanced in *snpc-1.2(-); snpc-1.3(-)* double mutants, suggesting that *snpc-1.2* functions partially redundantly with *snpc-1.3* to promote germline fertility. We next asked whether the loss of female piRNAs in *snpc-1.2(-)* mutants compromises fertility across generations. At 25°C, *prg-1(-)* mutants, like many mutants in germline small RNA pathways, exhibit either immediate sterility in the subsequent generation or a progressive germline mortal (Mrt) phenotype, characterized by a gradual decline in brood size over successive generations culminating in sterility (Buckley et al., 2012; Simon et al., 2014; Spracklin et al., 2017). In contrast, *snpc-1.2(-)* mutants remained fertile for at least 20 generations, although they consistently produced fewer and more variable numbers of progeny relative to wild type (Supplemental Fig. S4B). Together, these findings suggest that female piRNAs contribute to the maintenance of long-term germline health but are largely dispensable for fertility within a single generation.

### *snpc-1.2(-)* mutants do not exhibit detectable meiotic defects

We next examined whether the transgenerational subfertility in *snpc-1.2(-)* mutants results from meiotic defects during oogenesis. First, we analyzed chromosome axis and synaptonemal complex formation using HIM-3 and SYP-5 staining, respectively. Germlines dissected from *snpc-1.2(-)* and wild-type hermaphrodites grown at 25°C for one generation showed normal HIM-3 and SYP-5 localization, with proper colocalization along chromosomes (Supplemental Fig. S4C). To evaluate whether defects arise with prolonged temperature stress, we examined HIM-3 and SYP-5 in *snpc-1.2(-)* animals after three generations at 25°C. Even under these conditions, *snpc-1.2(-)* germlines displayed normal chromosome axis and synaptonemal complex formation, comparable to wild-type (Supplemental Fig. S4C).

We also tested for crossover defects by counting DAPI-stained bodies in oocytes at diakinesis. Normally, *C. elegans* oocytes display six DAPI-stained bodies, corresponding to six pairs of homologous chromosomes. We observed six DAPI bodies in both wild-type and *snpc-1.2(-)* oocytes even after three generations at 25°C, indicating normal crossover formation (Supplemental Fig. S4D). Finally, we assessed germline progression by quantifying mitotic, transition, and pachytene zones in dissected germlines. The *snpc-1.2(-)* mutants showed no significant differences in gross germline architecture compared to wild type across 1–3 generations at 25°C (Supplemental Fig. S4E). Together, these results suggest that the transgenerational subfertility observed in *snpc-1.2(-)* mutants is unlikely due to defects in meiosis.

### SNPC-1.2 and SNPC-1.3 collaborate to ensure proper development

Since *snpc-1.2(-)* mutants undergo normal meiosis, we next investigated whether they exhibit other physiological abnormalities. We first performed a high-incidence-of-males (Him) assay by quantifying the percentage of male progeny from P0 hermaphrodites grown at 25°C. While wild-type worms produced 0.3% male progeny, *prg-1(-)* worms produced 10.8% male progeny (Supplemental Fig. S4F). Single piRNA mutants *snpc-1.2(-)* and *snpc-1.3(-)* showed a modest increase (∼0.6%), while the *snpc-1.2(-); snpc-1.3(-)* double mutant displayed a synthetic Him phenotype, producing 3.1% male progeny (Supplemental Fig. S4F). These findings suggest that sex-specific piRNAs may contribute to accurate chromosome segregation by limiting chromosome nondisjunction.

We also examined developmental timing defects by measuring the time to first embryo laying in piRNA pathway mutants. Synchronized L1 larvae were monitored individually to determine when they laid their first embryo. The *snpc-1.2(-)* and *snpc-1.3(-)* single mutants showed normal developmental timing. In contrast, both *prg-1(-)* and *snpc-1.2(-); snpc-1.3(-)* double mutants exhibited a significant delay (Supplemental Fig. S4G). The observation that only the double mutant, but not either single mutant, displayed this developmental delay suggests that both male-specific and female-specific piRNAs are required together to promote timely germline development and reproductive output.

### SNPC-1.1 promotes snRNA but not piRNA expression

Previous mass spectrometry analysis of immunopurified SNPC-4 indicated that the SNPC-1 family member SNPC-1.1 interacts with SNPC-4 (Choi et al., 2021), prompting us to test whether SNPC-1.1, like SNPC-1.2 and SNPC-1.3, promotes piRNA expression. Unexpectedly, RNAi knockdown of *snpc-1.1* increased levels of both male and female piRNAs during spermatogenesis and oogenesis, respectively, indicating that SNPC-1.1 antagonizes rather than promotes piRNA expression (Supplemental Fig. S5A). These findings raised the possibility that SNPC-1.1 instead promotes an alternative noncoding RNA pathway and suppresses piRNA biogenesis by competing with SNPC-1.2 and SNPC-1.3 for access to SNPC-4. To test this, we examined whether SNPC-1.1 regulates snRNA expression. qPCR analysis revealed a significant reduction in U1 snRNA levels following *snpc-1.1* RNAi, comparable to the effect of *snpc-4* RNAi (Fig. 4A), supporting a role for SNPC-1.1 in snRNA expression.

**Figure 4.**
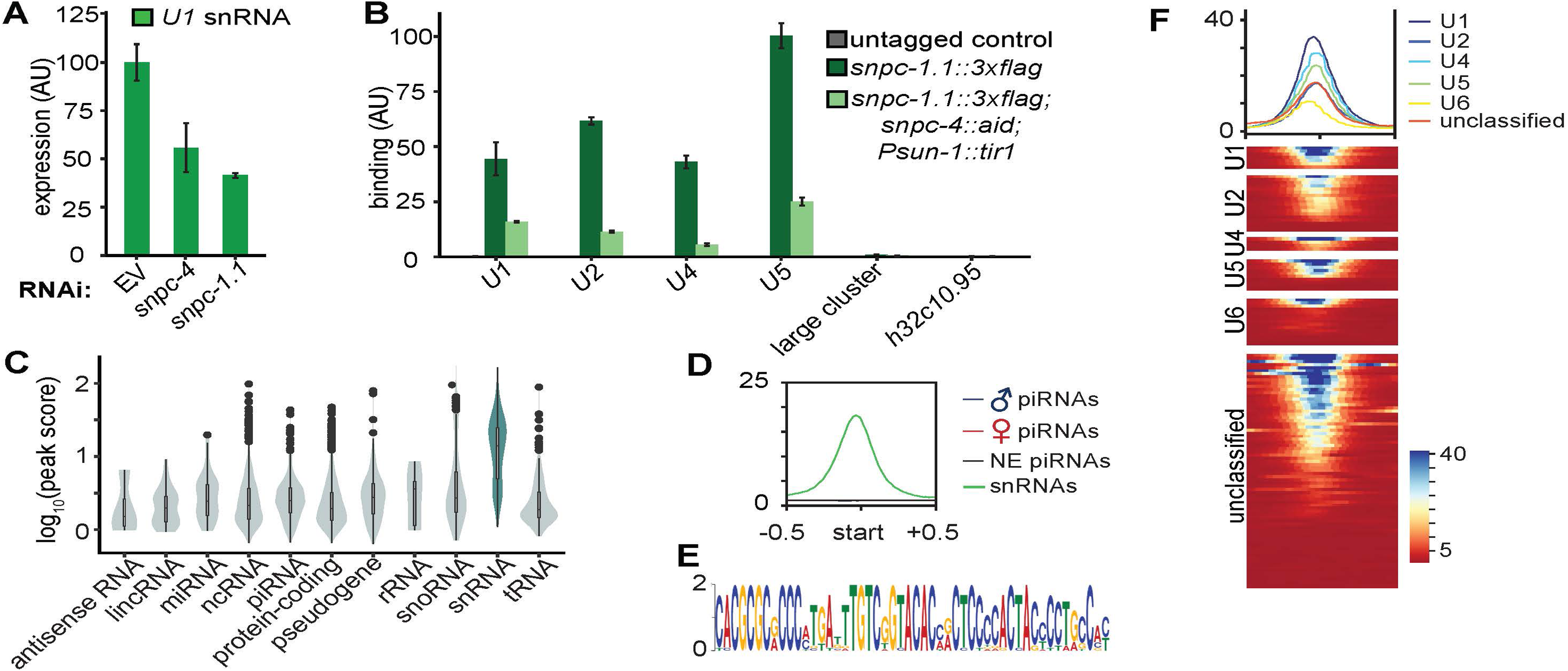
SNPC-1.1 binds snRNA loci and promotes snRNA transcription. **(A)** *snpc-1.1* knockdown reduces U1 snRNA levels. qPCR of U1 snRNA levels in RNAi-treated animals during oogenesis, normalized to *eft-2* mRNA. Error bars represent ± SD of two technical replicates. **(B)** SNPC-1.1 binds snRNA loci in a SNPC-4-dependent manner. SNPC-1.1::3xFlag ChIP-qPCR signal is normalized to input DNA (mean ± SD of two technical replicates) at U1, U2, U4, U5 snRNA genes; the large piRNA cluster on chromosome IV (13.5–17.2 Mb); and the noncoding gene *h32c10.95* on chromosome IV. SNPC-4::AID::Ollas was degraded upon auxin treatment (250 µM, 4 h). Error bars represent ± SD of two technical replicates. **(C)** SNPC-1.1 binding is enriched at snRNA loci. SNPC-1.1::3xFlag ChIP-seq binding profiles at various RNA gene classes. **(D)** SNPC-1.1 binds snRNA loci rather than piRNA loci. SNPC-1.1 ChIP-seq binding profiles at piRNA and snRNA loci. **(E)** Upstream sequences of SNPC-1.1–bound loci contain the *C. elegans* proximal sequence element (PSE), the canonical promoter element of snRNA genes. **(F)** SNPC-1.1 binds multiple snRNA classes. ChIP-seq binding profiles across snRNA classes.

To determine whether SNPC-1.1 directly associates with snRNA loci, we used CRISPR-Cas9 to endogenously tag SNPC-1.1 with a 3xFlag epitope and verified allele functionality by a fertility assay (Supplemental Fig. S5B). ChIP-qPCR using an untagged wild-type strain as a negative control revealed strong enrichment of SNPC-1.1::3xFlag at the U1, U2, U4, and U5 snRNA loci, but not at the chromosome IV large piRNA cluster or the non-coding gene *h32c10.95* (Fig. 4B). ChIP-seq at 56 hours post-L1 hatching confirmed genome-wide enrichment of SNPC-1.1 at snRNA genes (Fig. 4C). In contrast, SNPC-1.1 did not bind male, female, or NE piRNA loci (Fig. 4D) and instead associated with a distinct upstream sequence motif resembling the *C. elegans* proximal sequence element (PSE) characteristic of snRNA promoters (Thomas et al., 1990) (Fig. 4E). SNPC-1.1 was enriched across all major snRNA classes (Fig. 4F), and its chromatin association required SNPC-4, as auxin-mediated SNPC-4::AID depletion attenuated SNPC-1.1 ChIP signal (Fig. 4B). SNPC-1.1 protein levels were unchanged following SNPC-4 depletion (Supplemental Fig. S5C), suggesting that SNPC-4 promotes SNPC-1.1 chromatin recruitment but is dispensable for protein stability. Together, these findings support a model in which SNPC-1.1 directly drives snRNA transcription in an SNPC-4-dependent manner.

### SNPC-1.2 forms distinct nuclear foci from SNPC-1.1

Several piRNA transcription factors, including SNPC-4 and SNPC-1.3, localize to one or two nuclear foci in germ cells, likely corresponding to the two homologous piRNA clusters on chromosome IV (Ruby et al., 2006; Kasper et al., 2014; Weng et al., 2019; Weick et al., 2014; Choi et al., 2021). To assess whether SNPC-1.2 exhibits similar subnuclear localization, we performed immunofluorescence in *snpc-4::3xflag; snpc-1.2::ollas* worms. SNPC-1.2 formed discrete nuclear foci detected with an anti-Ollas antibody that colocalized with SNPC-4 during both spermatogenesis and oogenesis (Fig. 5B, D). No signal was detected in a *snpc-4::3xflag* control strain lacking the *snpc-1.2::ollas* tag (Fig. 5A, C), confirming specificity.

**Figure 5.**
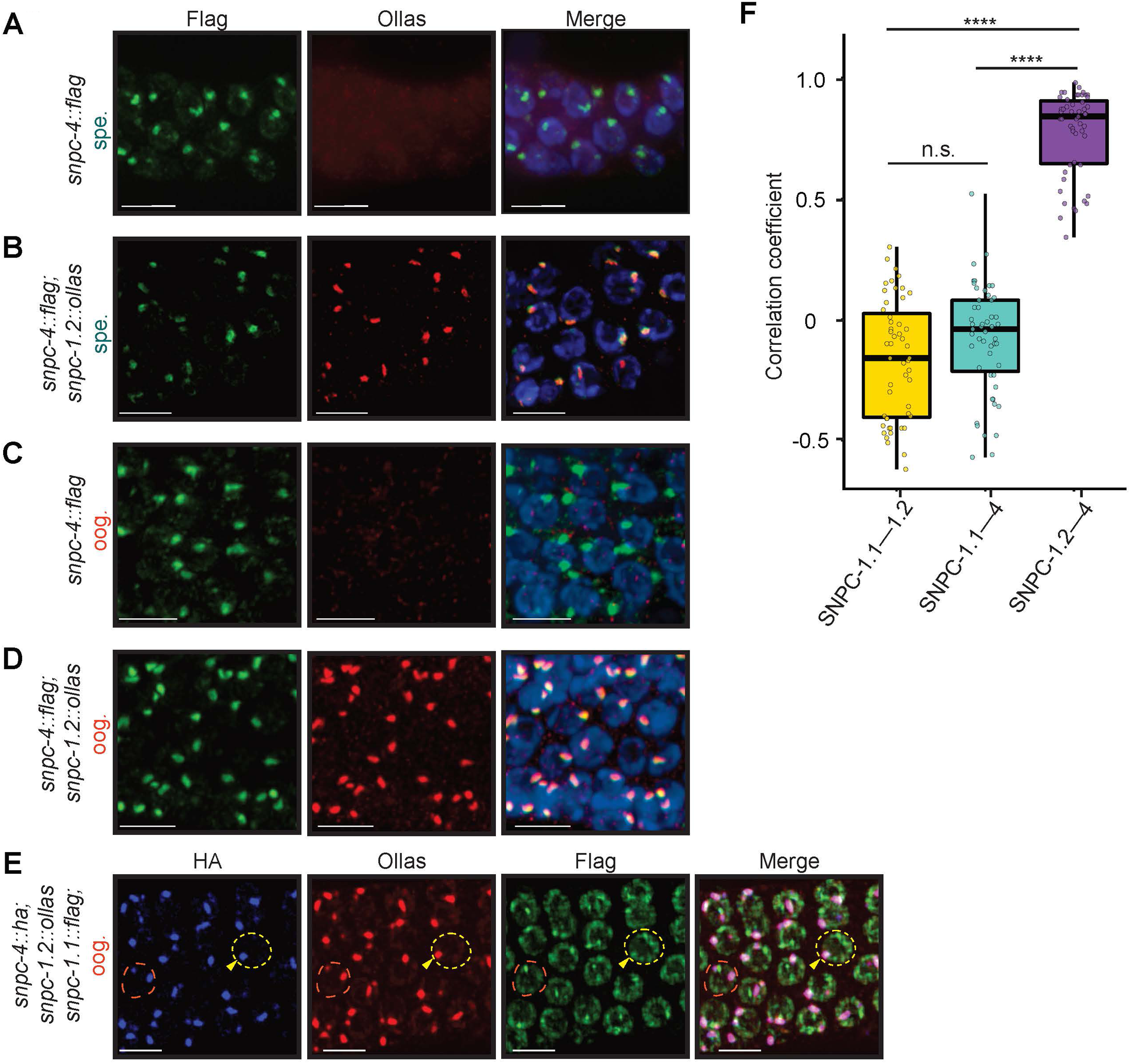
SNPC-1.2 forms foci with SNPC-4 that are adjacent to P granules and mostly distinct from SNPC-1.1 foci. **(A–D)** SNPC-1.2 forms foci with SNPC-4 in the germline. Dissected *snpc-4::3xflag* (first and third row) and *snpc-4::3xflag*; *snpc-1.2::ollas* (second and fourth row) germlines stained for SNPC-4::3xFlag (green), SNPC-1.2::Ollas (red), and DNA (blue) during spermatogenesis (spe.) and oogenesis (oog.). Representative images of meiotic (A) or mitotic (B–D) germ cells from three biological replicates are shown. Scale bar, 5 µm. **(E)** SNPC-1.2 foci overlap with SNPC-4 and are largely distinct from SNPC-1.1. Dissected *snpc-4::ha; snpc-1.2::ollas; snpc-1.1::3xflag* hermaphrodite germlines were stained for SNPC-4::HA (blue), SNPC-1.2::Ollas (red), and SNPC-1.1::3xFlag (green) during oogenesis. Representative image of meiotic germ cells from three biological replicates is shown. Yellow arrowhead indicates a focus with partial colocalization of SNPC-4, SNPC-1.2, and SNPC-1.1. Scale bar, 5 µm. **(F)** SNPC-1.2 colocalizes with SNPC-4 but not SNPC-1.1. Boxplots show Pearson correlation coefficients for SNPC-4::HA, SNPC-1.1::3xFlag, and SNPC-1.2::Ollas signals in meiotic nuclei from *snpc-4::ha; snpc-1.2::ollas; snpc-1.1::3xflag* germlines (n=50 nuclei). Statistical significance was assessed by Kruskal-Wallis H-test followed by Dunn’s test (n.s., non-significant; ****p<0.00005).

SNPC-1.1 and SNPC-1.2 regulate distinct small RNA classes: snRNA genes are dispersed across the genome, while piRNA genes are clustered. We therefore hypothesized that these transcription factors localize to distinct nuclear regions. To test this, we tagged SNPC-4 with HA at its C-terminus (*snpc-4::ha*) and crossed the allele into a *snpc-1.1::3xflag; snpc-1.2::ollas* strain. Immunofluorescence during oogenesis revealed that SNPC-1.2 and SNPC-4 colocalized in nuclear foci, whereas SNPC-1.1 displayed a more diffuse nucleoplasmic distribution. Notably, the brightest SNPC-1.1 signal did not overlap with the SNPC-1.2/SNPC-4 foci, although occasional weaker signals partially overlapped (Fig. 5E, F). These observations support spatial segregation of piRNA and snRNA transcriptional complexes and reinforce the functional specialization among SNPC-1 paralogs.

### Only SNPC-1.2 and SNPC-1.3 associate with piRNA-specific factors

To identify paralog-specific interaction partners, we performed immunoprecipitation followed by mass spectrometry in strains expressing 3xFlag-tagged SNPC-1.1, SNPC-1.2, or SNPC-1.3. SNPC-1.1 primarily interacted with SNPC-4 and TOFU-5 (Fig. 6A). Although TOFU-5 binds piRNA loci and associates with known piRNA factors (Weng et al., 2019), it has also been implicated in splice leader gene transcription during starvation, suggesting TOFU-5 has dual roles in snRNA and piRNA transcription like SNPC-4 (Hou et al., 2022). SNPC-1.1 additionally associated with SIP-1 and Y57A10A.2, implicating these proteins in snRNA transcription.

**Figure 6.**
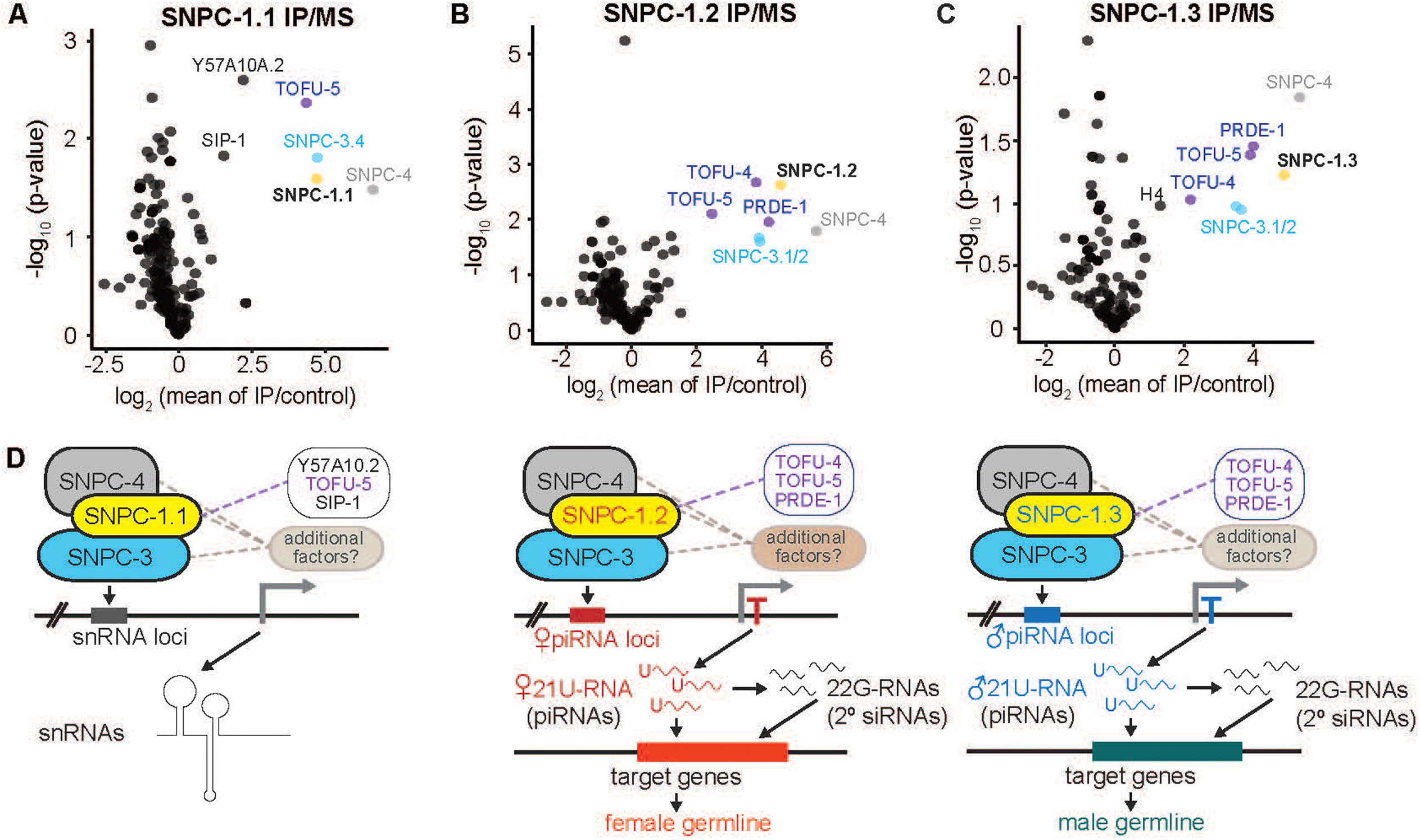
SNPC-1 paralogs specify snRNA, female piRNA, and male piRNA transcriptional complexes. **(A–C)** SNPC-1.2 and SNPC-1.3 IP/MS enriches piRNA transcription factors. Volcano plot showing enrichment in SNPC-1.1::3xFlag (left), SNPC-1.2 (middle), and SNPC-1.3 (right) immunoprecipitates relative to an untagged control, with significance values (n=3 biological replicates, n=2 for SNPC-1.3). Proteins exhibiting > 2-fold enrichment over the untagged control in all biological replicates are labeled. piRNA biogenesis factors (purple), SNPC-4 (gray), SNPC-1 (yellow), and SNPC-3 (blue) are indicated. **(D)** Model: SNPC-1 paralogs act as specificity factors for three distinct small-RNA transcriptional complexes. SNPC-1.1 associates with SNPC-4, SNPC-3.4, TOFU-5, and possibly other factors to promote snRNA transcription. In contrast, SNPC-1.2 assembles with SNPC-4 and other factors to constitute a female-specific USTC that promotes female piRNA transcription, whereas SNPC-1.3 assembles with SNPC-4 and other factors to constitute a male-specific USTC that promotes male piRNA transcription.

In contrast, SNPC-1.2 and SNPC-1.3 immunoprecipitations were enriched not only for SNPC-4 and TOFU-5, but also for the piRNA-specific factors PRDE-1 and TOFU-4 (Goh et al., 2014; Kasper et al., 2014; Weick et al., 2014; Weng et al., 2019) (Fig. 6B, C). Notably, none of the SNPC-1 proteins co-immunoprecipitated with other SNPC-1 paralogs, supporting a model in which SNPC-4 assembles into mutually exclusive transcriptional complexes, each containing a distinct SNPC-1 paralog and functioning within the larger USTC to promote sex-specific piRNA transcription (Fig. 6D). These findings suggest that duplication of the *snpc-1* gene family in *C. elegans* enabled functional diversification of an ancestral transcriptional complex, extending its role from snRNA transcription to sex-specific piRNA biogenesis.

To investigate whether expansion of *snpc-1* is unique to *C. elegans*, we performed a phylogenetic survey across the *Caenorhabditis* genus. Using Orthofinder, we identified *snpc-1.2* and *snpc-1.3* orthologs in other androdioecious (hermaphroditic) nematodes and in three of five dioecious (male/female) species (Supplemental Fig. S6A; Emms and Kelly, 2019). Broader analysis across nematode clades revealed that *snpc-1* duplication is largely restricted to clade V nematodes (Supplemental Fig. S6B), consistent with comparative epigenomics studies showing that the piRNA pathway is absent outside this clade (Beltran et al., 2019). These observations suggest that single-gene *snpc-1* orthologs in other nematodes likely function primarily in snRNA transcription.

## Discussion

### How is sex-specific piRNA and snRNA transcription coordinated?

While reuse of core machinery such as SNAPc in the transcription of multiple RNA classes is an efficient strategy, it raises the question of how each RNA class is precisely regulated throughout germline development. Our mass spectrometry analyses and previous structural studies support incorporation of a single SNPC-1 paralog into each SNAP complex (Fig. 6; Ma and Hernandez, 2002). We propose a model in which each SNAP complex includes a single, unique SNPC-1 paralog in a mutually exclusive manner, thereby directing transcription of distinct small RNA classes.

We also find that the expression level of each SNPC-1 paralog is crucial for determining the fidelity and timing of small RNA transcription. SNPC-1.3 is highly enriched during spermatogenesis (Choi et al., 2021), while SNPC-1.2 is preferentially expressed during oogenesis, likely reducing competition for SNPC-4 between these paralogs. However, in contexts where SNPC-1.2 and SNPC-1.3 are co-expressed, our data suggest they compete for shared transcriptional machinery. Female piRNAs are upregulated in *snpc-1.3(-)* mutants during spermatogenesis and downregulated when SNPC-1.3 is ectopically expressed during oogenesis (Fig. 1F; Choi et al., 2021). Conversely, male piRNAs increase modestly in *snpc-1.2(-)* mutants during oogenesis (Fig. 1F, Supplemental Fig. S1D), and piRNAs of both classes increase upon *snpc-1.1* RNAi (Supplemental Fig. S5A). These findings support a model in which SNPC-1 paralogs compete for limiting common components such as SNPC-4 and TOFU-5, thereby influencing transcription of their respective small RNA class. We hypothesize that coordination of SNPC-1 paralog expression, together with potential differences in affinity for core factors, ensures proper temporal and spatial expression of piRNAs and snRNAs.

### Functional modularity of the *C. elegans* SNAP complexes

Our findings show that SNAPc retains its ancestral role in snRNA transcription via SNPC-1.1, while SNPC-1.2, SNPC-1.3, and possibly SNPC-1.4 have acquired new functions in piRNA transcription. While SNPC-1.2 and SNPC-1.3 have distinct sex-specific roles, it is possible that SNPC-1.4 may have a spatiotemporal role—such as in pre-pachytene or pachytene piRNA transcription—although this needs to be tested. We propose that the modular assembly of SNAPc, comprising three interacting subunits, facilitated this co-option, enabling the emergence of novel functions without compromising essential snRNA production. Similar modular repurposing has been observed for PETISCO, whose function changes from piRNA precursor processing or embryogenesis depending on alternative adaptor proteins PID-1 or TOST-1 (Cordeiro Rodrigues et al., 2019; Zeng et al., 2019; Wang et al., 2021). These examples suggest that co-option of ancestral RNA-processing complexes facilitated evolution of the piRNA pathway.

### How do SNPC-1 paralogs specify the transcription of the correct RNA class?

SNPC-4 in *C. elegans* plays an expanded role compared to its homologs, contributing to the transcription of snRNAs, snoRNAs, tRNAs, and piRNAs (Kasper et al., 2014). It is plausible that different SNPC-1 paralogs direct SNPC-4 to specific classes of RNA genes. While the precise mechanism remains unclear, work in flies has shown that all SNAPc subunits are required for binding to the proximal sequence element (PSE) at snRNA genes, and each SNAPc subunit can directly bind DNA (Henry et al., 1996; Wong et al., 1998; Li et al., 2004; Hung and Stumph, 2012). However, recent cryo-EM studies of human SNAPc suggest that only SNAPC3 and SNAPC4 make direct contact with DNA, with SNAPC1 functioning in a regulatory or structural capacity (Sun et al., 2022), suggesting possible divergence in subunit function across species.

Our ChIP-seq data show that SNPC-4 binds male piRNA genes even in the absence of *snpc-1.3*, but fails to bind female piRNA genes in *snpc-1.2(-)* mutants (Fig. 3G, Supplemental Fig. S3C, D). This suggests that SNPC-1.3 may activate transcription at loci where SNPC-4 is already bound, whereas SNPC-1.2 may be required for SNPC-4 recruitment to female piRNA loci. One potential reason for SNPC-4 binding male piRNAs in *snpc-1.3(-)* is that SNPC-1.2 compensates for SNPC-1.3 and guides the complex to male piRNAs. Consistent with this, SNPC-1.2 is expressed during spermatogenesis (Fig. 1C), binds male piRNA loci (Fig. 3E), and partially compensates for male piRNA loss in *snpc-1.3(-)* mutants (Fig. 1F). Examining SNPC-4 association with male piRNA genes in *snpc-1.2(-); snpc-1.3(-)* background would test this possibility.

We propose two models to explain how SNPC-1 paralogs confer transcriptional specificity: 1) SNPC-4 binds target loci independently, and SNPC-1 recruitment initiates transcription through complex-specific cofactors; or 2) SNPC-1 binds SNPC-4 prior to DNA association and guides the complex to specific target promoters using distinct DNA-binding features. Our observation that *snpc-1.3(-)* mutants retain SNPC-4 binding but lose male piRNA expression supports a model in which SNPC-1.3 functions primarily in transcriptional activation. In contrast, the near-complete loss of SNPC-4 binding at female piRNA genes in *snpc-1.2(-)* mutants suggests a recruitment role.

The recent identification of ATTF-6, an AT-hook transcription factor required for USTC recruitment to piRNA clusters (Wang et al., 2025), suggests that piRNA locus recognition involves multiple DNA-binding factors. ATTF-6 may help establish the initial chromatin context that allows SNPC-4 to access piRNA promoters, either independently (as we observe for male piRNAs) or in an SNPC-1.2-dependent manner (as we observe for female piRNAs). Additionally, the role of chromatin remodeling in exposing piRNA promoters (Paniagua et al., 2024) and the formation of phase-separated transcriptional condensates at piRNA clusters (Zhu et al., 2025) indicate that SNPC-1 paralog specificity operates within a highly organized nuclear environment. Understanding how SNPC-1 paralogs integrate with these architectural features will be important for fully elucidating the mechanisms of sex-specific piRNA transcription. Future work dissecting the SNPC-1 protein domains that confer specificity will help distinguish between these models.

### What are the functions of female piRNAs?

Although SNPC-1.2 is required for transcription of a major subclass of piRNAs, *snpc-1.2(-)* mutants do not display pronounced fertility defects (Supplemental Fig. S4A) (Cox et al., 1998; Batista et al., 2008). In contrast, *snpc-1.3(-)* mutants exhibit severe single-generation fertility defects, similar to those observed in *prg-1(-)* null animals (Choi et al., 2021; Shi et al., 2013). Male piRNA targets are depleted for spermatogenesis genes (Shi et al., 2013). Recent work has shown that piRNAs initiate transcriptional silencing of hundreds of spermatogenic genes during spermatogenesis, directly promoting sperm differentiation and function (Cornes et al., 2022). Consistent with this, loss of male piRNAs in *snpc-1.3(-)* mutants leads to defective spermatogenesis (Choi et al., 2021). These findings suggest that fertility defects in *prg-1(-)* mutants are driven primarily by the loss of male piRNAs. In contrast, the gradual fertility decline observed in *snpc-1.2(-)* mutants over multiple generations (Supplemental Fig. S4B) raises the possibility that female piRNAs may contribute to germline maintenance over time.

The distinct phenotypes of *snpc-1.2(-)* and *snpc-1.3(-)* mutants suggest that, while male and female piRNAs likely cooperate in conserved functions such as transposon silencing, they also exhibit sex-specific functional specialization. Male piRNAs may regulate gene expression required for spermatogenesis, whereas female piRNAs may mediate genome surveillance. The *C. elegans* piRNA pathway initiates RNA-induced epigenetic silencing (RNAe) including single-copy transgenes like *gfp::cdk-1,* which are silenced in wild type but expressed in *prg-1(-)* mutants (Shirayama et al., 2012). These findings suggest that female piRNAs may serve a broader role in identifying and buffering against foreign sequences, while male piRNAs may have more specific targets required for proper spermatogenesis.

Manipulating *snpc-1.2* and *snpc-1.3* enables a relatively easy, albeit incomplete, separation of male and female piRNA expression, offering a powerful system for dissecting sex-specific piRNA function. Additionally, our identification of SNPC-1.1 as a dedicated snRNA transcription factor establishes the *C. elegans snpc-1* gene family as a model for studying how transcriptional complexes acquire specificity. Overall, our findings reveal how an ancestral complex, SNAPc, has diversified in *C. elegans* to coordinate the expression of distinct classes of noncoding RNAs.

## Materials and Methods

### Strain generation

Strains generated by CRISPR/Cas9 genome editing were produced as described previously (Paix et al., 2015). crRNAs and repair templates are provided in Supplemental Table 8; strains used in this study are listed in Supplemental Table 9.

### Quantitative RT-PCR

cDNA synthesis prior to qPCR analysis was performed as described previously (Weiser et al., 2017; Choi et al., 2021). For *him-8(-)* qPCR assays, approximately 450 animals were picked into M9 buffer. Primer sequences are provided in Supplemental Table 8.

### RNAi Assays

Bacterial RNAi clones were obtained from the Ahringer RNAi library (Kamath et al., 2003) and experiments performed as previously described (Choi et al., 2021)

### Chromatin immunoprecipitation

Liquid culture growth at 20°C was performed as described previously (Zanin et al., 2011). Approximately 2.5 million animals were grown per 2-L flask in 500 mL complete S Basal medium. 250µM auxin was added to *snpc-1.2::3xflag; snpc-4::aid::ollas; Psun-1::TIR1* and *snpc-1.1::3xflag; snpc-4::aid::ollas; Psun-1::TIR1* worms before collection. For SNPC-1.2 ChIP, animals were cleaned by sucrose flotation prior to live cross-linking. For staged collections at 48 and 72 h post-L1 hatching, animals were harvested at 48 h and remaining animals were transferred to fresh complete S Basal containing HB101. For collection, the gut was cleared for 15 min while nutating in M9, followed by three washes in M9. Cross-linking was performed as described in Choi et al., 2021. For SNPC-1.1, germline nuclei were isolated as detailed in Kirshner et al., 2025. ChIP was performed as described previously (Weiser et al., 2017), with the addition of proteinase K during de-crosslinking. Extracted DNA was resuspended in 18 µL 1 M Tris-EDTA pH 8 and 2 µL of 20 mg/mL RNase A (Invitrogen).

### ChIP-qPCR

ChIP-qPCR reactions were performed in 25-μL volumes containing 2 μL ChIP DNA and SYBR Green master mix (Thermo Fisher) with 35 nM forward and reverse primers. Reactions were run on a Bio-Rad CF96 Real-Time PCR system and normalized to input.

### Covalent crosslinking of Dynabeads

Protein G Dynabeads (Thermo Fisher) were conjugated to monoclonal mouse anti-FLAG M2 antibody (Sigma-Aldrich) as described in Choi et al., 2021

### Immunoprecipitation

Immunoprecipitation was performed as described previously (Choi et al., 2021).

### Western blotting

Prior to gel electrophoresis, dithiothreitol (DTT) was added to samples to a final concentration of 0.1 M, and samples were heated at 95°C for 15 min. Western blotting protocol and antibody dilutions are delineated in Choi et al., 2021.

### Mass spectrometry and analysis

Mass spectrometry was performed at the Johns Hopkins Mass Spectrometry and Proteomics Facility. For analysis, a value of one was added to total spectral counts to avoid zeros. Spectral counts for each protein were normalized to total spectral counts within each immunoprecipitation sample. Fold enrichment was calculated by dividing normalized spectral counts from the tagged strain by those from the untagged control for each biological replicate and then averaging across replicates. Enrichment values were expressed as log_2_ fold-change, and statistical significance was subsequently calculated.

### Immunofluorescence microscopy

Synchronized L4 larvae or adult animals were dissected and stained as described in Choi et al., 2021. Primary antibodies used were mouse M2 anti-Flag antibody (1:200; Sigma-Aldrich), rabbit anti-HA (1:1,000; Cell Signaling Technology), and rat anti-Ollas (1:100; Novus Biologicals). For staining of meiotic germline samples, rabbit anti-SYP-5 (1:500) and chicken anti-HIM-3 (1:500) antibodies were provided by the Yumi Kim laboratory (Johns Hopkins University).

Secondary antibodies used were Alexa-Fluor 488 anti-mouse (1:300), Alexa-Fluor 647 anti-rabbit (1:500), Alexa-Fluor 555 anti-rat (1:300), and Alexa-Fluor anti-chicken (1:300). Images from three biological replicates were acquired using a Zeiss LSM 700 confocal microscope or Leica Thunder Imager. Representative images were processed by generating maximum intensity projection from z-stacks using FIJI or Leica software.

### Germline mortality assay

Germline mortality assays were performed as described previously (Weiser et al., 2017).

### Hermaphrodite fertility assays

To assess single-generation fertility, gravid animals were bleached and embryos plated onto NGM plates seeded with OP50 at either 20°C or 25°C as indicated. At the L2 or L3 stage, animals were singled to individual plates, and the progeny were counted.

### Mating assays

To assess potential mating defects, P_0_ hermaphrodites of both mating strains were maintained at 20°C and synchronized by hypochlorite treatment. Synchronized L1 larvae were plated onto NGM plates seeded with OP50 at 25°C and allowed to develop. Prior to the L4 stage, five males and one female were transferred to a new plate at 25°C. To ensure that all F_1_ progeny were cross-progeny, we crossed the *fem-1(hc17)* allele into the background of all hermaphrodite strains. At 25°C, *fem-1(hc17)* causes complete feminization of the germline. F_1_ progeny were scored at the L4 stage or later to assess fertility.

### Time-to-embryo-lay assay

Hermaphrodites of each genotype were synchronized by hypochlorite treatment, and embryos (P_0_) were hatched overnight in M9 buffer. Synchronized L1 larvae were plated onto NGM plates seeded with OP50 at either 20°C or 25°C and allowed to develop to the L2/L3 stage before being singled to individual plates. Starting at 40 h post-hatch at 25°C, P_0_ animals were examined hourly to determine the onset of embryo laying.

### Him assay

To assess male progeny production, hermaphrodites were synchronized by hypochlorite treatment, and synchronized L1 larvae were plated onto NGM plates seeded with OP50 at 25°C. At the L2/L3 stage, 3–5 P_0_ animals of each genotype were transferred to individual plates at 25°C.

At the L4 stage, plates were inspected to confirm that no males had been transferred. F_1_ progeny were scored starting at the L4 stage or later, and the incidence of males was quantified.

### Quantitative and statistical analysis

Unless otherwise indicated, all quantitative analyses are shown as mean ± standard deviation (SD), with SD shown as error bars. Two independent biological replicates were performed for germline mortality assays and co-immunoprecipitation experiments. Three independent biological replicates were performed for all other experiments, with one representative replicate shown.

### Small RNA-seq library preparation, sequencing, and analysis

One of the small RNA-seq datasets used in this study was published previously (Choi et al., 2021). Three additional datasets were generated at the High-Throughput Sequencing Core at the University of California, Los Angeles. Briefly, 2 µg total RNA was pretreated with RNA 5’ polyphosphatase (Illumina) as described previously (Phillips et al., 2014). Pretreated RNA was used for small RNA library preparation with the NEBNext Multiplex Small RNA Library Prep kit (New England BioLabs) according to the manufacturer’s instructions.

One set of libraries was sequenced as 75-bp single-end reads, and two sets were sequenced as 150-bp paired-end reads on a NovaSeq 6000 platform (Illumina). For paired-end libraries, only R1 reads trimmed to 50 bp were used for downstream analysis. Raw reads were trimmed for Illumina adapters and quality using Trimmomatic v0.39 (SLIDING WINDOW: 4:25) (Bolger et al., 2014). Reads between 19 and 30 nt in length were retained. Reads 21–23 nt in length were defined as piRNAs and aligned to the *C. elegans* reference genome (WBcel235) using Bowtie v1.1.1 with parameters -v 0 k 5 –best –strata –tryhard (Langmead et al., 2009). piRNAs were quantified using a custom Python-based pipeline built upon the HTSeq framework. Read assignments were performed using the intersection-nonempty overlap resolution mode with the nonunique=fraction fractional counting setting. Putative novel piRNAs were identified from reads 21 nt in length by incorporating additional 5’ positional and nucleotide matching criteria to detect piRNAs absent from the reference annotation. Putative novel piRNAs were then filtered against the ncRNA reference annotation to exclude reads corresponding to other annotated ncRNAs and were further defined based on expression support across wild-type samples, requiring a minimum raw read count of ≥5 per sample in at least half of the wild-type samples (n = 6) at either the spermatogenic or oogenic timepoint.

Quality control of raw and aligned reads was performed using FastQC v0.11.7 (http://www.bioinformatics.babraham.ac.uk/projects/fastqc/) and SAMtools 1.9 (Li et al., 2009). Principal component analysis was conducted on raw count data following normalization with edgeR (Robinson et al., 2010).

To identify differentially expressed piRNAs, an expression filter of counts per million (CPM) > 2 in at least 3 samples was applied to remove lowly expressed piRNAs. Differential expression analysis was performed using edgeR, comparing mutants to wild type, with thresholds of log_2_(fold-change) ≥ 0.26 and false discovery rate (FDR) ≤ 0.05 (Benjamini–Hochberg). To account for batch effects across four datasets, an additive model including batch was incorporated into the design matrix. For motif discovery, nucleotide sequences corresponding to the 60 nt upstream of each piRNA locus were extracted from the reference genome and analyzed using MEME Suite v5.1.1 (Bailey et al., 2009).

### ChIP-seq library preparation, sequencing, and analysis

ChIP-seq libraries were prepared and multiplexed using the Ovation Ultralow Library Systems v2 kit (NuGEN Technologies) according to the manufacturer’s protocol. Libraries were sequenced as 50-bp paired-end reads on a NovaSeq 6000 platform (Illumina). Demultiplexed raw data were processed as described previously (Choi et al., 2021).

Global enrichment of SNPC-1.2 ChIP-seq signal at piRNA clusters on chromosome IV was assessed by calculating normalized read coverage across consecutive, non-overlapping 1-kb bins across the *C. elegans* genome.

Binding profiles and heatmaps for SNPC-1.1, SNPC-1.2, SNPC-1.3, and SNPC-4 at piRNA and snRNA loci were generated using MACS v2.1.2 (Zhang et al., 2008) and deepTools (Ramírez et al., 2016). Fold-enrichment signal tracks were produced from read count-normalized genome-wide pileup and lambda track outputs using callpeak (bdgcmp in MACS2). Enrichment signal track regions overlapping with regions in the ENCODE ce 11 blacklist (https://github.com/Boyle-Lab/Blacklist/) were removed from further analysis. Using bigwigCompare within deepTools, enrichment signal tracks in bigWig format were then normalized (ratio) to the corresponding no-tag control. Binding profiles and heatmaps were then made using deepTools (computeMatrix followed by plotHeatmap).

### Orthology assignment

To identify orthology relationships, proteomes were obtained from WormBase Parasite (https://parasite.wormbase.org/) and analyzed using OrthoFinder (Emms and Kelly, 2019). Relevant gene trees for orthogroups containing the *snpc-1* gene family were visualized and processed using Interactive Tree of Life v6 (iTOL) (Letunic and Bork, 2021).

## Data and software availability

Newly generated small RNA-seq and ChIP-seq data have been deposited in the NCBI Gene Expression Omnibus (GEO) under accession numbers GSE341621 and GSE341964, respectively. This study also incorporates previously generated data (GEO accession GSE152831; Choi et al., 2021). Code used for the analyses presented in this study is available on Github (https://github.com/starostikm/SNPCs).

## Acknowledgments

We thank the members of the Kim Lab (Charlotte Choi, Himani Galagali, Greg Fuller, Amelia Alessi, Jessie Kirshner, Anna Vakhnovetsky, Youngyong Park, Victoria Murphy, Haena Lee, Tian Tan, Yuqi Tang, Julia Zhou, and Daniel Lenchner) for their insightful recommendations on this project. We thank Bob Cole and Raquel Shortt of the Johns Hopkins University Mass Spectrometry and Proteomics Core for their help with the mass spectrometry experiments. The *Caenorhabditis* Genetics Center provided several strains used in this project. This work was supported by the National Institutes of Health (R01HD109667 to J.K.K, F31HD108916 to L.K.B, and F31HD107960 to M.R.S.). S.E.J. is an Investigator of the Howard Hughes Medical Institute.

## Author contributions

L.K.B., M.R.S., R.J.T., M.C.S., and J.K.K. conceptualized the project. L.K.B., M.R.S., R.J.T., M.C., M.C.S., and J.K.K. designed experiments and interpreted the results. L.K.B., M.R.S., R.J.T., A.I., M.C. performed the experiments. S.F. and S.E.J. generated the sequencing data. M.R.S. performed the bioinformatics analyses. L.K.B., M.R.S., R.J.T., and J.K.K. wrote and edited the manuscript. J.K.K. oversaw the project.

---

**Supplemental Figure 1.**
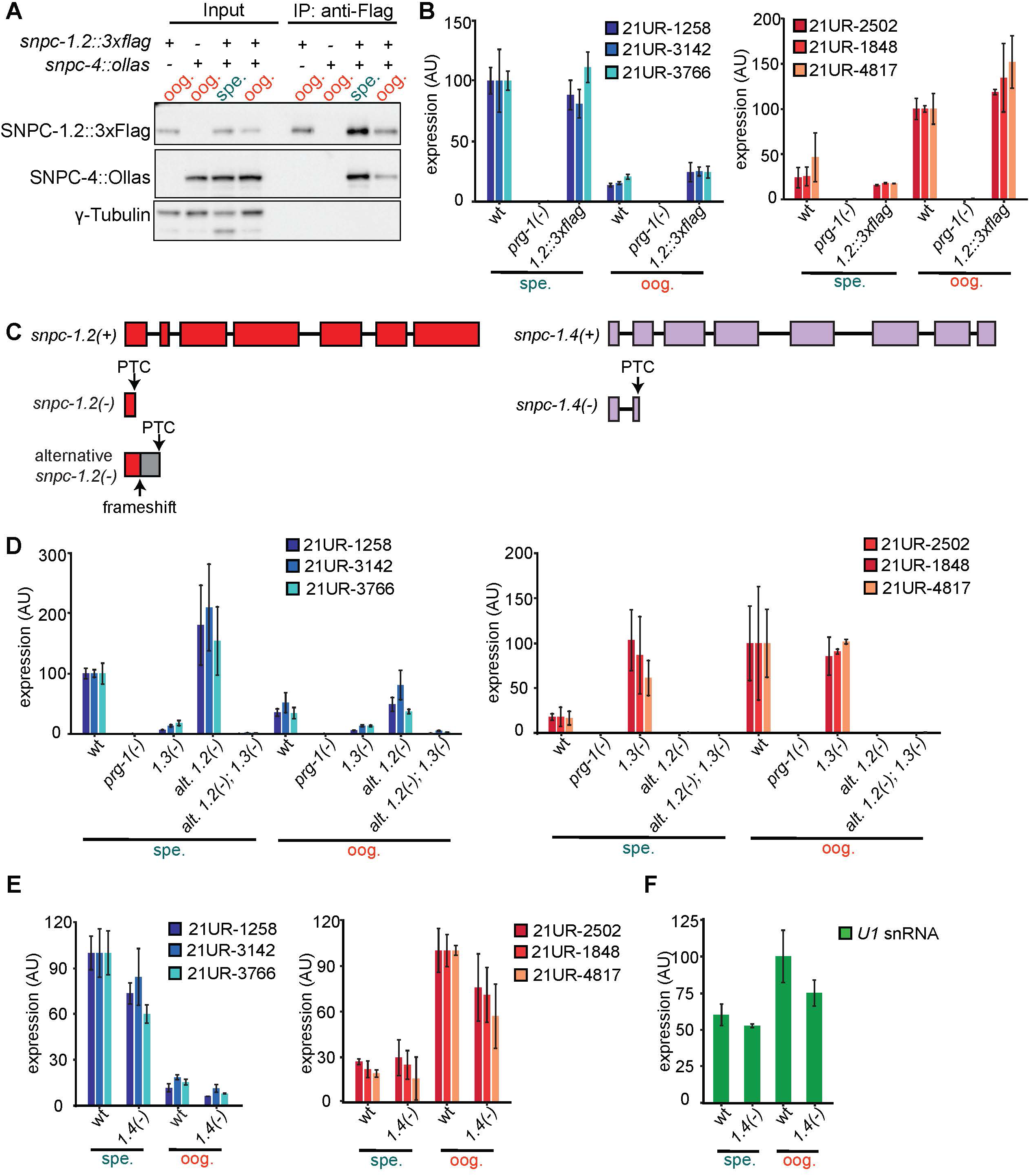
SNPC-1.2, but not SNPC-1.4, is required for female piRNA transcription. **(A)** SNPC-1.2 interacts with SNPC-4. Anti-Flag immunoprecipitation of SNPC-1.2::3xFlag during spermatogenesis and oogenesis. γ-tubulin serves as a loading control. **(B)** SNPC-1.2::3xFlag retains wild-type function. TaqMan qPCR of male (left) and female (right) piRNA levels during spermatogenesis (spe.) and oogenesis (oog.) in the *snpc-1.2::3xflag* strain. Expression is normalized to U18 snoRNA. Error bars represent ± SD of two technical replicates. **(C)** Schematic of wild-type and mutant *snpc-1.2* (left) and *snpc-1.4* (right) alleles used in this study. PTC, premature termination codon. **(D)** An independent *snpc-1.2* deletion allele shows reduced female piRNA expression. TaqMan qPCR of male (left) and female (right) piRNAs during spermatogenesis (spe.) and oogenesis (oog.) normalized to U18 snoRNA. Error bars represent ± SD of two technical replicates. **(E)** SNPC-1.4 modestly promotes piRNA expression. TaqMan qPCR of male (left) and female (right) piRNAs during spermatogenesis (spe.) and oogenesis (oog.) normalized to U18 snoRNA. Error bars represent ± SD of two technical replicates. **(F)** SNPC-1.4 does not markedly alter U1 snRNA levels. TaqMan qPCR of U1 snRNA during spermatogenesis (spe.) and oogenesis (oog.) normalized to *eft-2* mRNA. Error bars represent ± SD of two technical replicates.

**Supplemental Figure 2.**
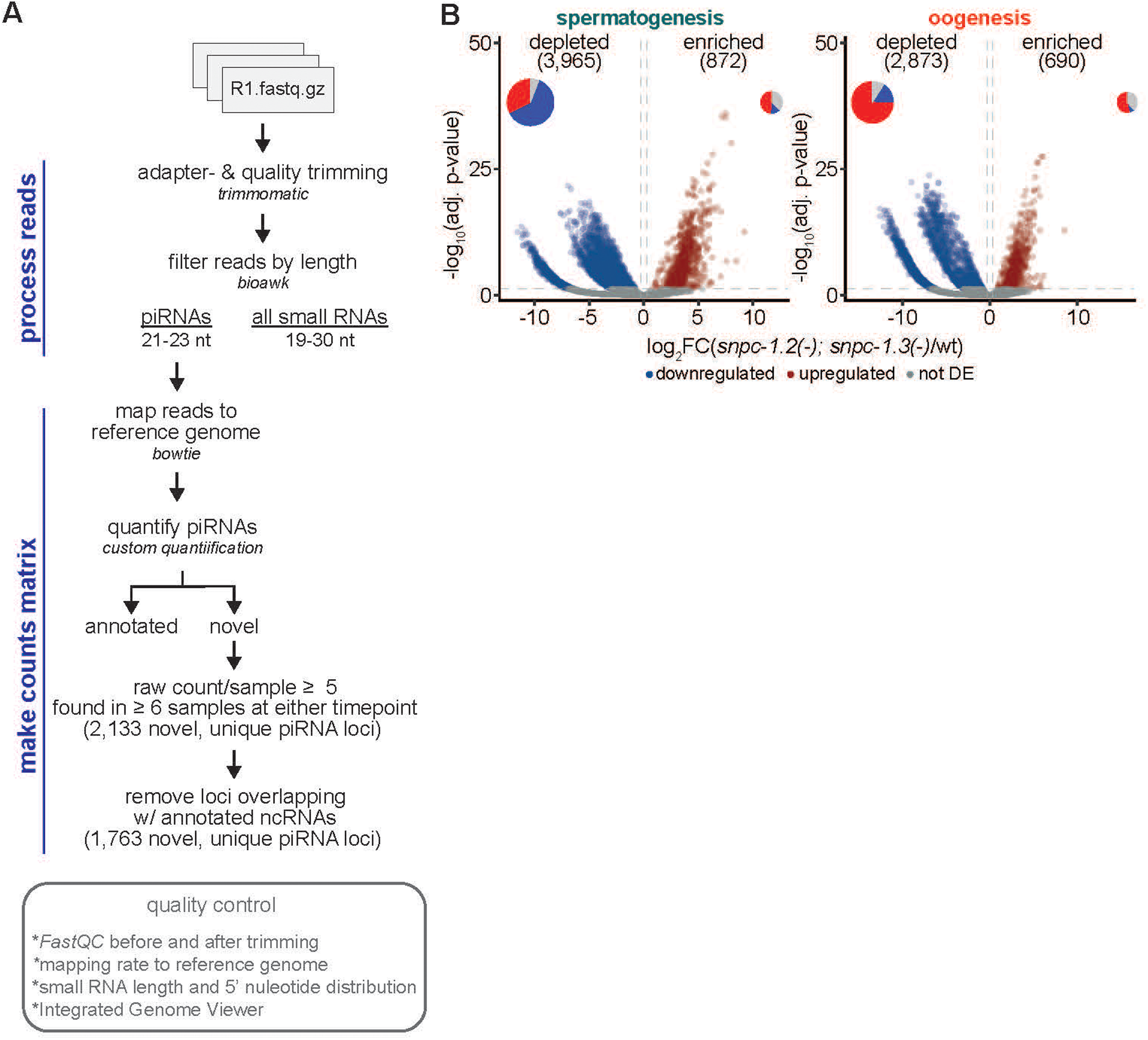
Small RNA-seq analysis workflow. **(A)** Annotated and novel piRNAs were identified from 21–23-nt reads that mapped to the WBcel235 reference genome and quantified using our custom read-counting pipeline. **(B)** piRNAs are globally down-regulated in *snpc-1.2(-); snpc-1.3(-)* double mutants. Volcano plots show significant down-regulation of male piRNAs during spermatogenesis (left) and male and female piRNAs during oogenesis (right).

**Supplemental Figure 3.**
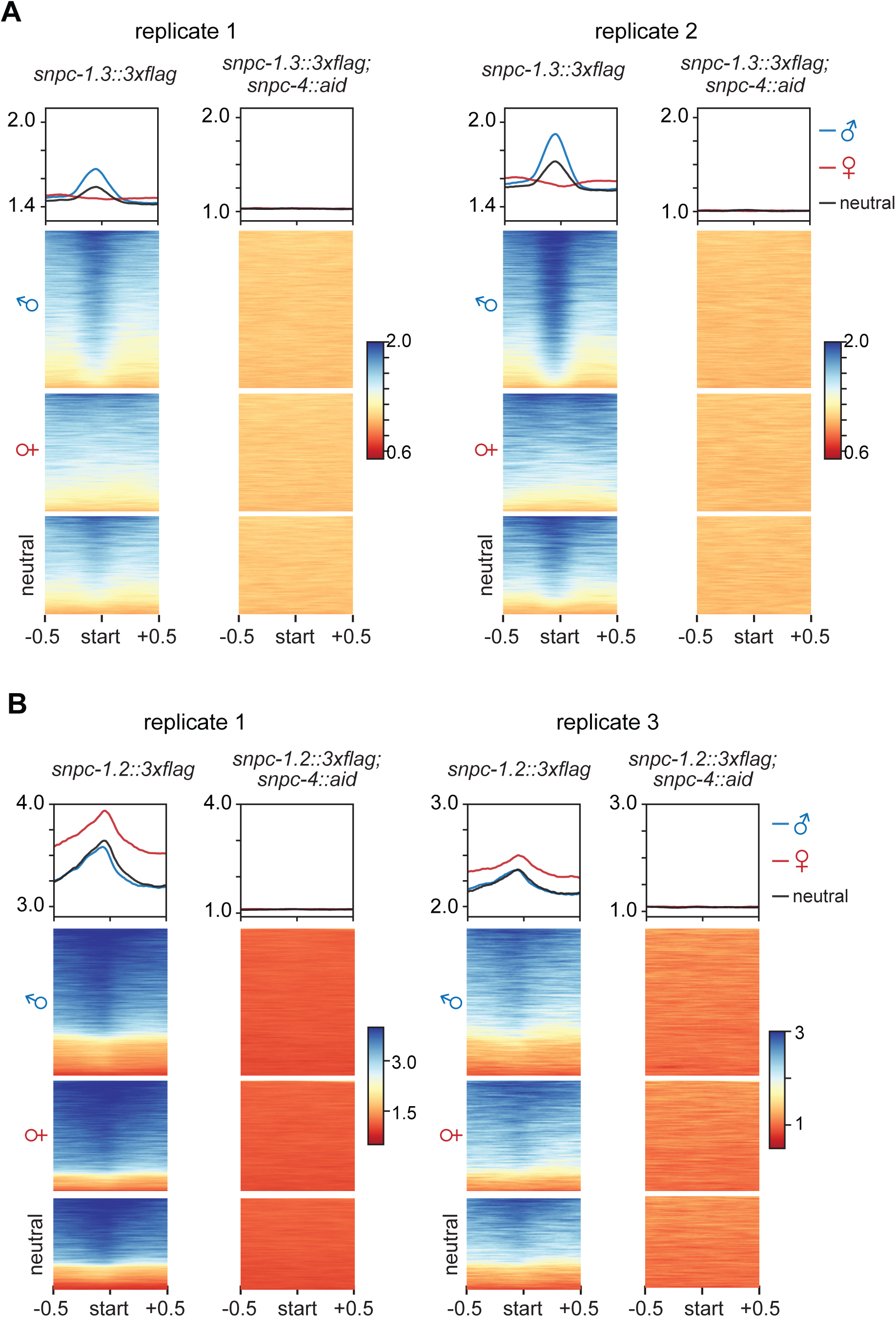
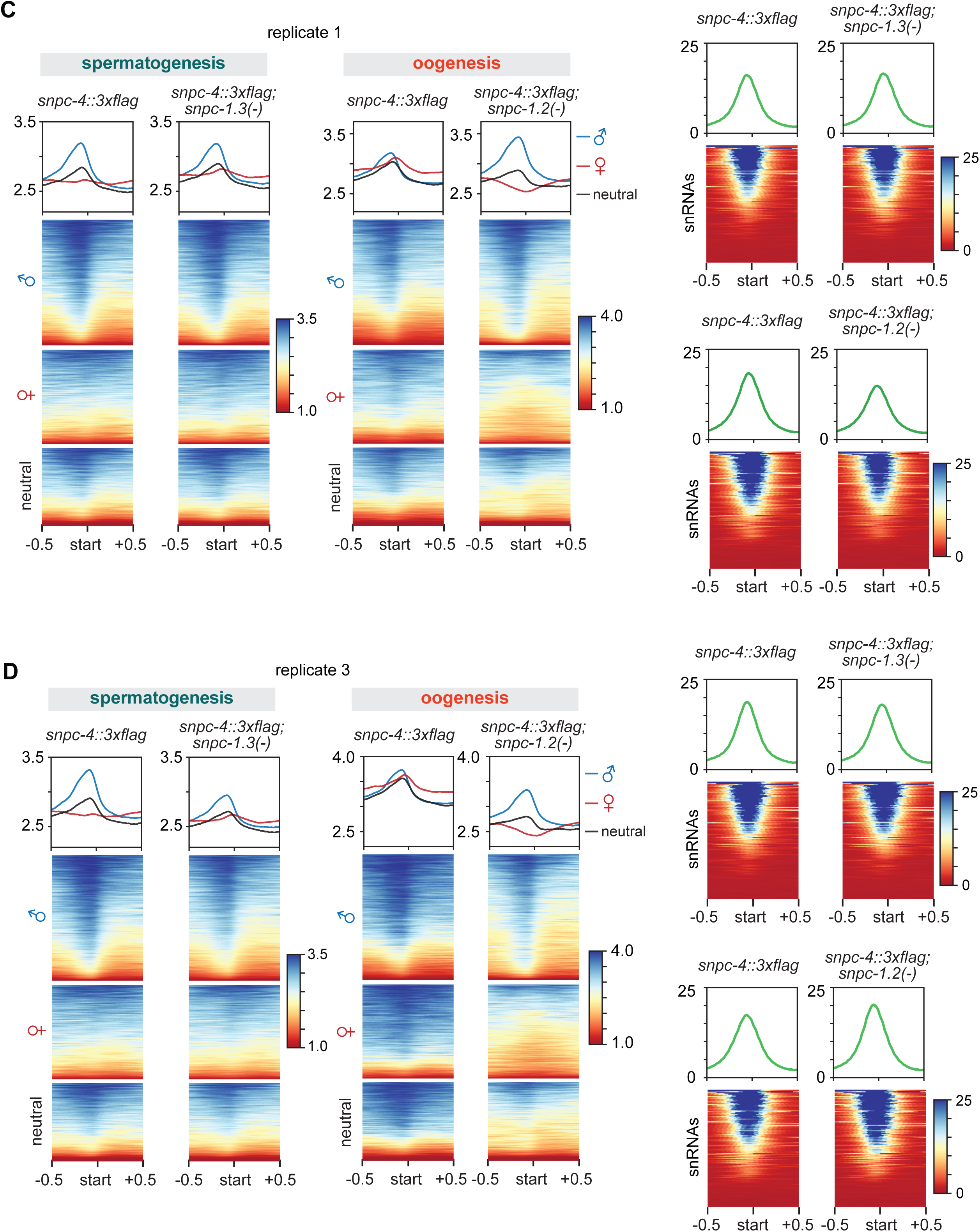
SNPC-1 piRNA factors bind piRNA loci in a *snpc-4*-dependent manner. **(A)** SNPC-4 is required for SNPC-1.3 binding to piRNA loci. Two biological replicates show SNPC-1.3 read distribution (mean density ± SE) centered on the 5′ nucleotide of mature piRNAs. Heat maps represent ChIP signal at the same sites. **(B)** SNPC-4 is required for SNPC-1.2 enrichment to female piRNA loci. Two biological replicates show SNPC-1.2 read distribution (mean density ± SE) centered on the 5′ nucleotide of mature piRNAs. Heat maps represent ChIP signal at the same sites. **(C,D)** SNPC-4 requires *snpc-1.2* for female piRNA locus binding but not *snpc-1.3* for male locus binding. Left: Two biological replicates showing SNPC-4 read distribution (mean density ± SE) centered on the 5′ nucleotide of mature piRNAs in wild-type, *snpc-1.3(-),* and *snpc-1.2(-)* backgrounds. Heat maps show ChIP signal at the same sites. Right: Two biological replicates showing SNPC-4 read distribution (mean density ± SE) centered on the 5’ nucleotide of snRNA genes in the same genotypes; heat maps show ChIP signal centered the 5′ nucleotide of snRNAs.

**Supplemental Figure 4.**
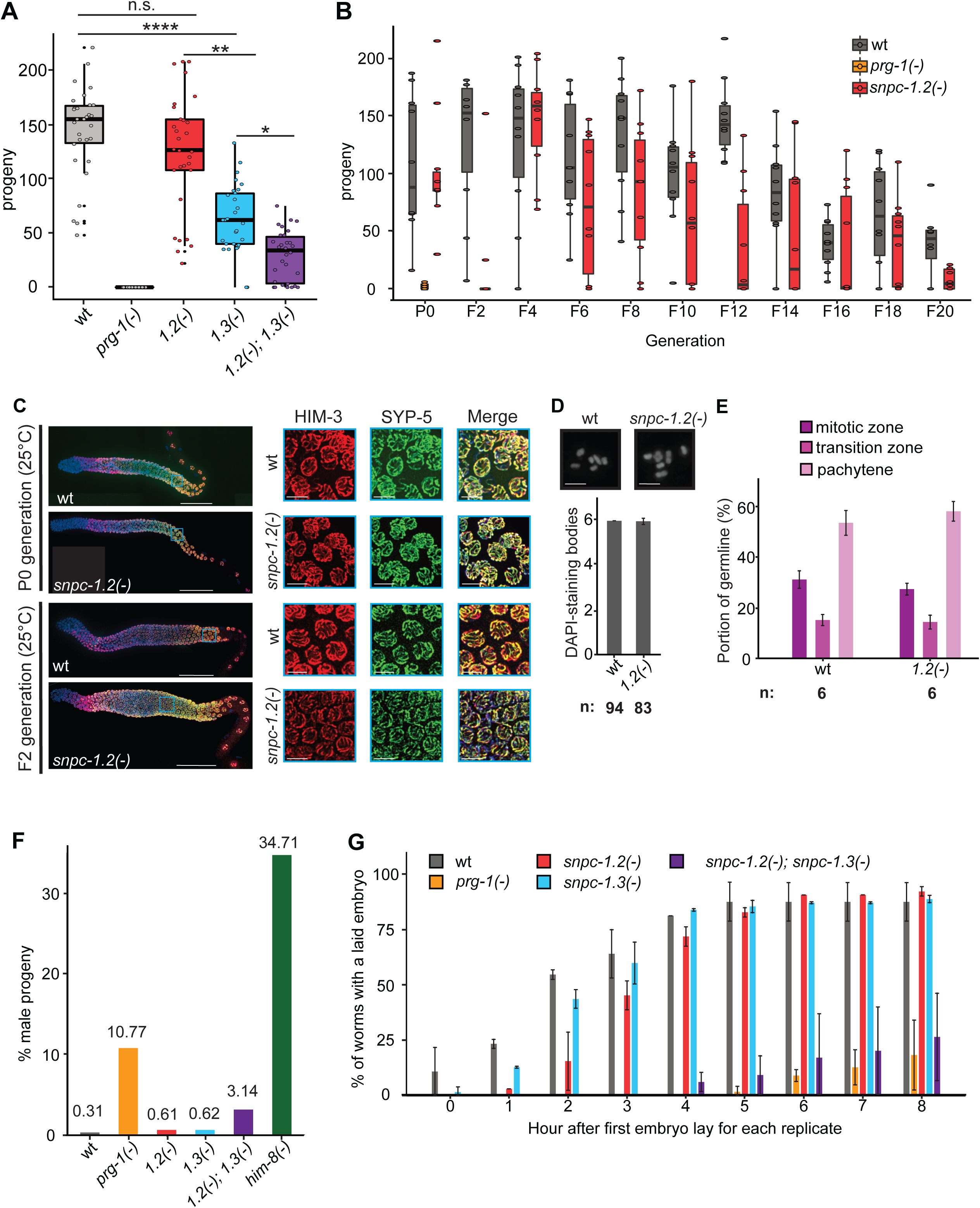
s*n*pc*-1.2(-)* mutants show transgenerational subfertility that is not due to meiotic defects during oogenesis. **(A)** *snpc-1.2* does not reduce fertility in a wild-type background. Each point represents viable progeny from a single hermaphrodite (n=30). Single-generation fertility assay conducted at 25°C. Boxes represent the 25^th^–75^th^ percentiles (Q1–Q3). The interquartile range (IQR) is defined as Q3–Q1. The center line indicates the median, and whiskers extend to the most extreme values within 1.5xIQR. Statistical significance was assessed by Kruskal-Wallis H-test followed by Dunn’s test (n.s., non-significant; *p<0.05, **p<0.005, ****p<0.00005). **(B)** *snpc-1.2* modestly promotes fertility over generations. Each point represents the progeny from a single worm (n=10 per genotype). Transgenerational fertility assay conducted at 25°C. Box-and-whisker plots are displayed as in (A). **(C)** *snpc-1.2(-)* mutants show normal chromosomal axis and synaptonemal complex formation during meiosis. Left: Wild-type and *snpc-1.2(-)* hermaphrodites were grown at 25°C for either one (P_0_) or three generations (F_2_) and germlines were dissected and stained for DNA (blue), axis marker HIM-3 (red), and synaptonemal complex protein SYP-5 (green). Scale bar, 50 µm. Right: Insets show pachytene nuclei. Representative images from three biological replicates are shown. Scale bar, 5 µm. **(D)** *snpc-1.2(-)* mutants show normal crossover formation. Top: Wild-type and *snpc-1.2(-)* oocytes at diakinesis stained with DAPI. Bottom: Quantification of DAPI-stained bodies per oocyte; total oocytes counted are indicated. Animals were maintained at 25°C for one, two, or three generations. Error bars represent ± SD of three biological replicates. Scale bar, 5 µm. **(E)** *snpc-1.2(-)* mutants progress normally through meiosis. Germline regions corresponding to mitotic and meiotic stages were quantified after one, two, or three generations at 25°C. Two germlines were analyzed per generation (six germlines total per genotype). Rows of nuclei were counted by DAPI staining and expressed as a fraction of total rows through diakinesis. Error bars represent ± SD of three biological replicates. **(F)** *snpc-1.2(-); snpc-1.3(-)* double mutants produce increased male progeny. Him assay performed at 25°C. Percentage of F1 male progeny shown atop the bars. Total worms counted for the six genotypes listed from left to right are n=8837, 7485, 10016, 10847, 5631, 3212. **(G)** *snpc-1.2* and *snpc-1.3* promote the proper timing of embryo laying. Time-to-embryo-lay assay conducted at 25°C. The y-axis indicates the percentage of animals that laid an embryo. Percentages were calculated from two biological replicates (32–35 P_0_ animals per replicate). The x-axis indicates hours after the first embryo lay in each replicate. Error bars represent ± SD of two biological replicates.

**Supplemental Figure 5.**
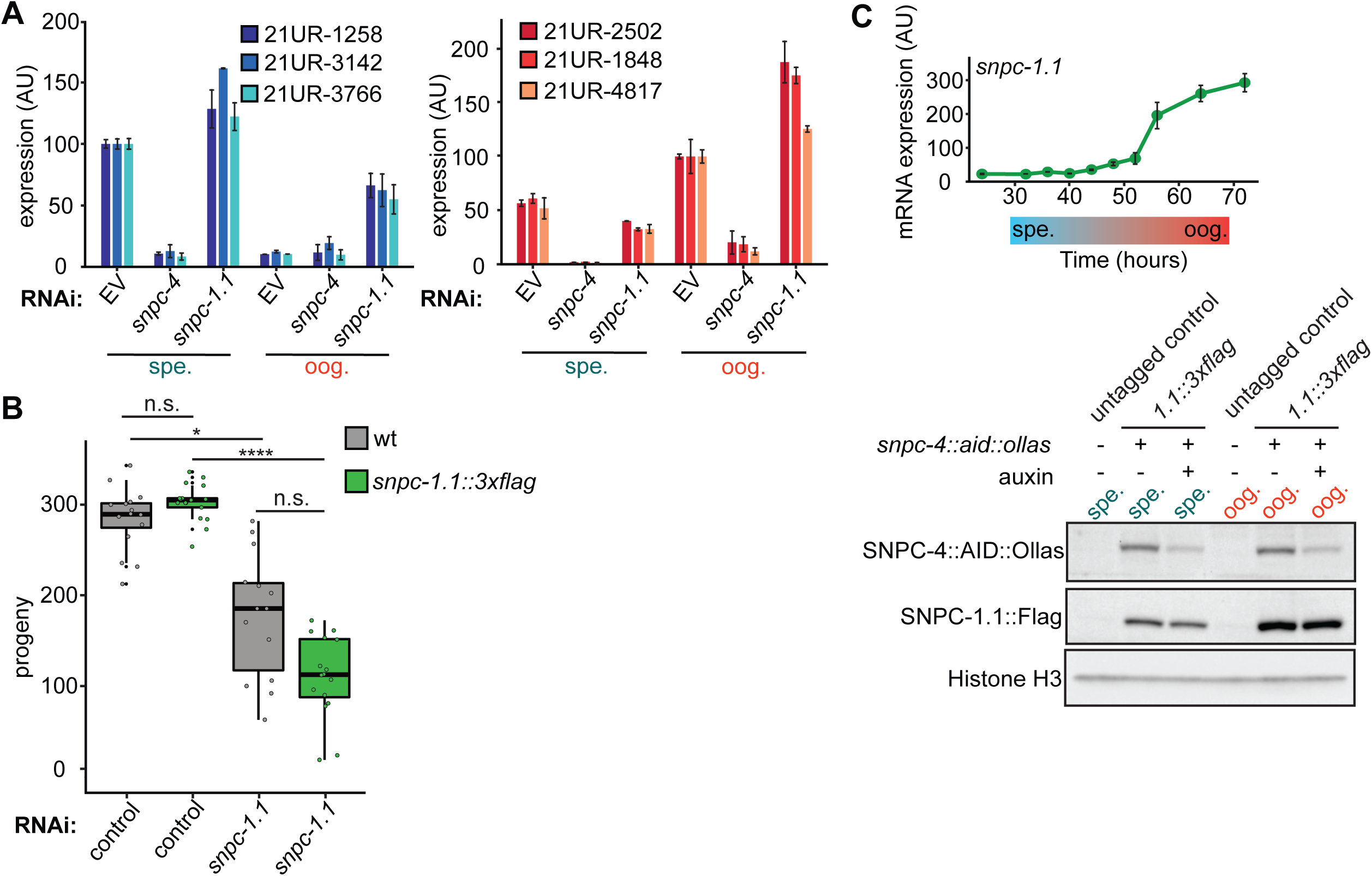
SNPC-1.1 is not required for piRNA expression. **(A)** SNPC-1.1 is not required for male or female piRNA expression. TaqMan qPCR of male (left) and female (right) piRNAs at spermatogenesis (spe.) and oogenesis (oog.). Expression normalized to U18 snoRNA. Error bars represent ± SD of two technical replicates. **(B)** The *3xflag* tag at the endogenous *snpc-1.1* locus does not impair function. Single-generation brood assay of wild-type and *snpc-1.1::3xflag* animals fed *gfp* RNAi (negative control) or *snpc-1.1* RNAi at 20°C (n=14–17). Each point represents viable progeny from a single hermaphrodite. Boxes indicate the interquartile range (IQR) with the center line marking the median; whiskers extend to 1.5xIQR. Statistical significance was assessed by Kruskal-Wallis H-test followed by Dunn’s test (n.s., non-significant; *p<0.05, ****p<0.00005). **(C)** *snpc-1.1* mRNA is elevated during oogenesis, and SNPC-1.1 protein levels are not dependent on SNPC-4. Top: qPCR time-course of *snpc-1.1* mRNA expression across development, normalized to *eft-2.* Error bars represent ± SD of two technical replicates. Bottom: Western blot of SNPC-1.1::3xFlag during spermatogenesis and oogenesis following SNPC-4::AID::Ollas depletion with auxin (250 µM, 4 h). Histone H3 serves as a loading control.

**Supplemental Figure 6.**
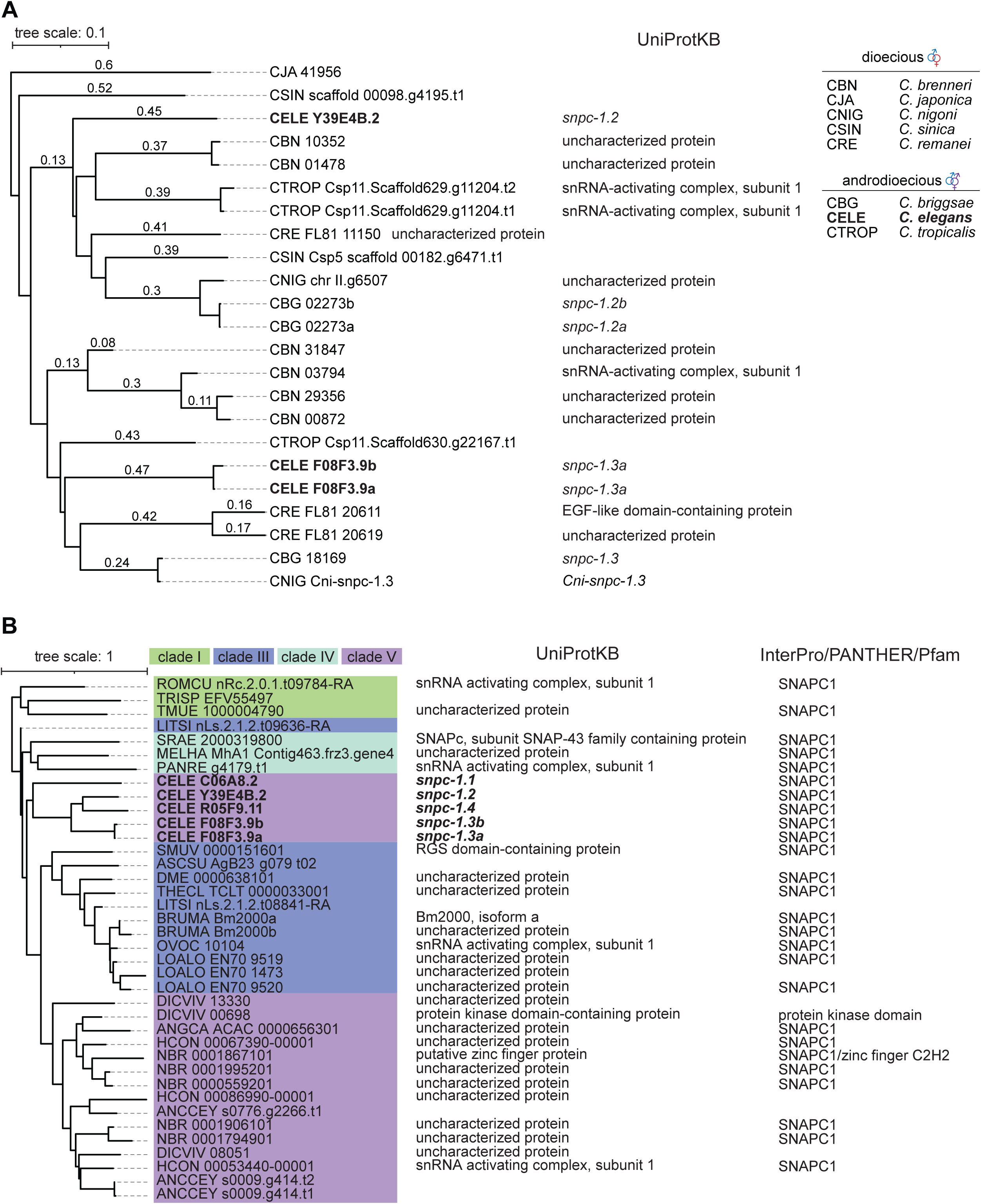
Orthologs of the *C. elegans snpc-1* gene family. **(A)** Conservation of sex-specific piRNA biogenesis factors in dioecious and androdioecious nematodes. Left: OrthoFinder identification of orthologs of *C. elegans snpc-1.2* and *snpc-1.3*. Right: associated UniProtKB gene names. **(B)** *snpc-1* gene duplication is restricted to clade V nematodes. Left: OrthoFinder analysis with nematode clades highlighted (clade I, green; clade III, blue; clade IV, light blue; clade V, purple). Right: associated UniProtKB gene names and protein domain annotations from InterPro, PANTHER, and Pfam.

**Supplemental Figure 7.**
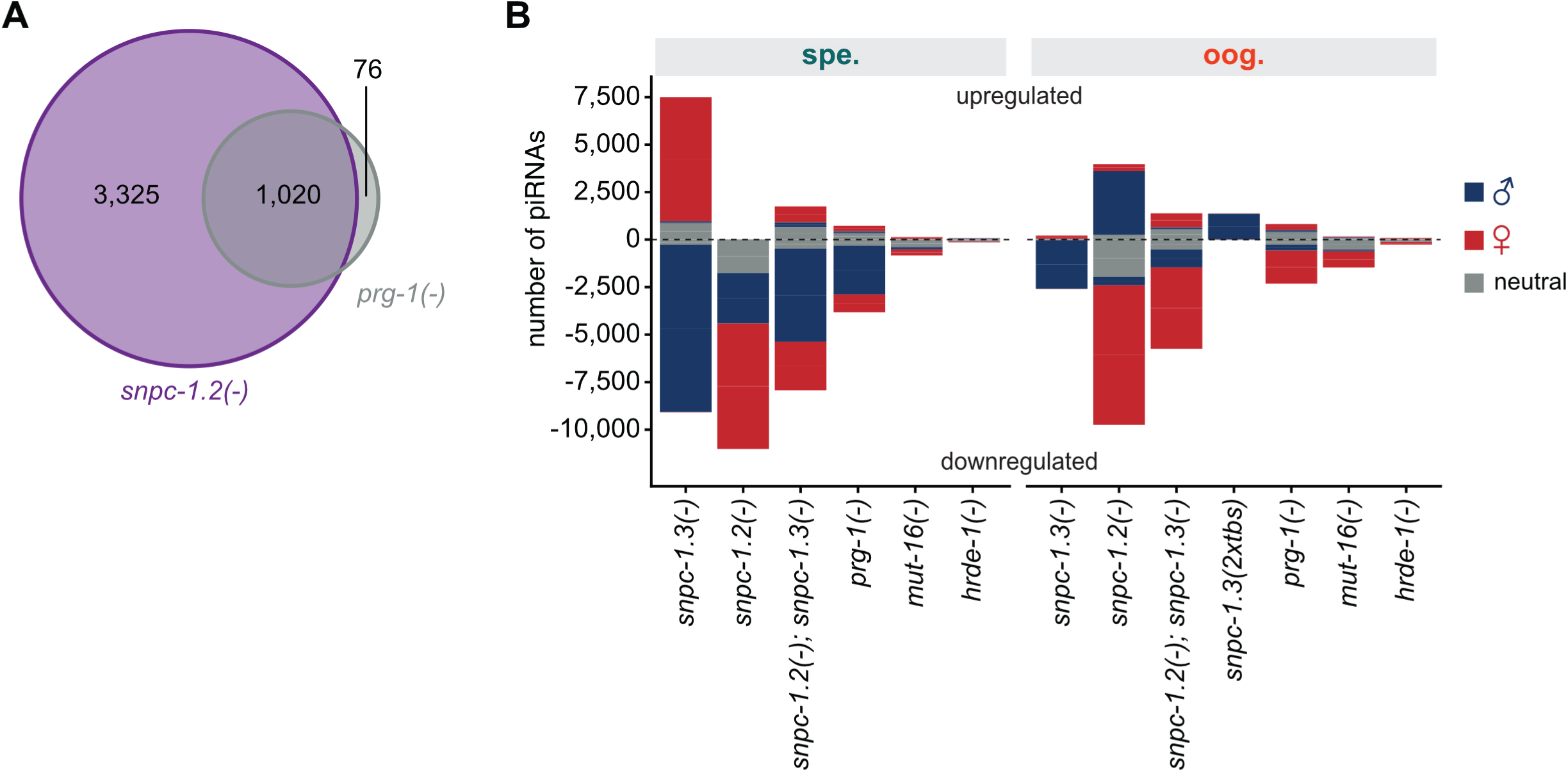
Overlap and sex-specific effects of defects in small RNA pathways. **(A)** piRNAs depleted in *prg-1(-)* mutants are also depleted in *snpc-1.2(-)* mutants. Venn diagram showing the overlap between piRNAs significantly depleted in *prg-1(-)* and *snpc-1.2(-)* mutants during oogenesis. **(B)** Sex-specific effects of piRNA and other small RNA pathway mutants on piRNA expression. Summary of differential expression analyses showing the numbers of male, female, and neutral (or not enriched) piRNAs significantly up- or downregulated in the indicated mutants relative to wild type during spermatogenesis or oogenesis.

## Notes

### Competing Interest Statement

The authors have declared no competing interest.

## References

Anders S, Pyl PT, Huber W. 2015. HTSeq--a Python framework to work with high-throughput sequencing data. Bioinformatics 31:166–169.

Aravin AA, Sachidanandam R, Girard A, Fejes-Toth K, Hannon GJ. 2007. Developmentally regulated piRNA clusters implicate MILI in transposon control. Science 316:744–747.

Bailey TL, Boden M, Buske FA, Frith M, Grant CE, Clementi L, Ren J, Li WW, Noble WS. 2009. MEME SUITE: tools for motif discovery and searching. Nucleic Acids Res 37:W202–8.

Batista PJ, Ruby JG, Claycomb JM, Chiang R, Fahlgren N, Kasschau KD, Chaves DA, Gu W, Vasale JJ, Duan S et al. 2008. PRG-1 and 21U-RNAs interact to form the piRNA complex required for fertility in *C. elegans*. Mol Cell 31:67–78.

Beanan MJ, Strome S. 1992. Characterization of a germ-line proliferation mutation in *C. elegans*. Development 116:755–766.

Beltran T, Barroso C, Birkle TY, Stevens L, Schwartz HT, Sternberg PW, Fradin H, Gunsalus K, Piano F, Sharma G et al. 2019. Comparative epigenomics reveals that RNA polymerase II pausing and chromatin domain organization control nematode piRNA biogenesis. Dev Cell 48:793–810.e6.

Billi AC, Freeberg MA, Day AM, Chun SY, Khivansara V, Kim JK. 2013. A conserved upstream motif orchestrates autonomous, germline-enriched expression of *Caenorhabditis elegans* piRNAs. PLoS Genet 9:e1003392.

Bolger AM, Lohse M, Usadel B. 2014. Trimmomatic: a flexible trimmer for Illumina sequence data. Bioinformatics 30:2114–2120.

Brennecke J, Aravin AA, Stark A, Dus M, Kellis M, Sachidanandam R, Hannon GJ. 2007. Discrete small RNA-generating loci as master regulators of transposon activity in *Drosophila*. Cell 128:1089–1103.

Buckley BA, Burkhart KB, Gu SG, Spracklin G, Kershner A, Fritz H, Kimble J, Fire A, Kennedy S. 2012. A nuclear Argonaute promotes multigenerational epigenetic inheritance and germline immortality. Nature 489:447–451.

Chen P, Kotov AA, Godneeva BK, Bazylev SS, Olenina LV, Aravin AA. 2021. piRNA-mediated gene regulation and adaptation to sex-specific transposon expression in *D. melanogaster* male germline. Genes Dev 35:914–935.

Choi CP, Tay RJ, Starostik MR, Feng S, Moresco JJ, Montgomery BE, Xu E, Hammonds MA, Schatz MC, Montgomery TA et al. 2021. SNPC-1.3 is a sex-specific transcription factor that drives male piRNA expression in *C. elegans*. Elife 10. doi:10.7554/eLife.60681

Cordeiro Rodrigues RJ, de Jesus Domingues AM, Hellmann S, Dietz S, de Albuquerque BFM, Renz C, Ulrich HD, Sarkies P, Butter F, Ketting RF. 2019. PETISCO is a novel protein complex required for 21U RNA biogenesis and embryonic viability. Genes Dev 33:857–870.

Cornes E, Bourdon L, Singh M, Mueller F, Quarato P, Wernersson E, Bienko M, Li B, Cecere G. 2022. piRNAs initiate transcriptional silencing of spermatogenic genes during *C. elegans* germline development. Dev Cell 57:180–196.e7.

Cox DN, Chao A, Baker J, Chang L, Qiao D, Lin H. 1998. A novel class of evolutionarily conserved genes defined by piwi are essential for stem cell self-renewal. Genes Dev 12:3715– 3727.

de Albuquerque BFM, Luteijn MJ, Cordeiro Rodrigues RJ, van Bergeijk P, Waaijers S, Kaaij LJT, Klein H, Boxem M, Ketting RF. 2014. PID-1 is a novel factor that operates during 21U-RNA biogenesis in *Caenorhabditis elegans*. Genes Dev 28:683–688.

Deng W, Lin H. 2002. miwi, a murine homolog of piwi, encodes a cytoplasmic protein essential for spermatogenesis. Dev Cell 2:819–830.

Doniach T, Hodgkin J. 1984. A sex-determining gene, *fem-1*, required for both male and hermaphrodite development in *Caenorhabditis elegans*. Dev Biol 106:223–235.

Emms DM, Kelly S. 2019. OrthoFinder: phylogenetic orthology inference for comparative genomics. Genome Biol 20:238.

Gasior SL, Wakeman TP, Xu B, Deininger PL. 2006. The human LINE-1 retrotransposon creates DNA double-strand breaks. J Mol Biol 357:1383–1393.

Girard A, Sachidanandam R, Hannon GJ, Carmell MA. 2006. A germline-specific class of small RNAs binds mammalian Piwi proteins. Nature 442:199–202.

Goh W-SS, Seah JWE, Harrison EJ, Chen C, Hammell CM, Hannon GJ. 2014. A genome-wide RNAi screen identifies factors required for distinct stages of *C. elegans* piRNA biogenesis. Genes Dev 28:797–807.

Henry RW, Ma B, Sadowski CL, Kobayashi R, Hernandez N. 1996. Cloning and characterization of SNAP50, a subunit of the snRNA-activating protein complex SNAPc. EMBO J 15:7129–7136.

Hinkley CS, Hirsch HA, Gu L, LaMere B, Henry RW. 2003. The small nuclear RNA-activating protein 190 Myb DNA binding domain stimulates TATA box-binding protein-TATA box recognition. J Biol Chem 278:18649–18657.

Hodgkin J, Horvitz HR, Brenner S. 1979. Nondisjunction mutants of the nematode *Caenorhabditis elegans*. Genetics 91:67–94.

Hou X, Zhu C, Xu M, Chen X, Sun C, Nashan B, Guang S, Feng X. 2022. The SNAPc complex mediates starvation-induced trans-splicing in *Caenorhabditis elegans*. J Genet Genomics 49:952–964.

Houwing S, Kamminga LM, Berezikov E, Cronembold D, Girard A, van den Elst H, Filippov DV, Blaser H, Raz E, Moens CB et al. 2007. A role for Piwi and piRNAs in germ cell maintenance and transposon silencing in Zebrafish. Cell 129:69–82.

Hung K-H, Stumph WE. 2011. Regulation of snRNA gene expression by the *Drosophila melanogaster* small nuclear RNA activating protein complex (DmSNAPc). Crit Rev Biochem Mol Biol 46:11–26.

Hung K-H, Stumph WE. 2012. Localization of residues in a novel DNA-binding domain of DmSNAP43 required for DmSNAPc DNA-binding activity. FEBS Lett 586:841–846.

Hung K-H, Titus M, Chiang S-C, Stumph WE. 2009. A map of *Drosophila melanogaster* small nuclear RNA-activating protein complex (DmSNAPc) domains involved in subunit assembly and DNA binding. J Biol Chem 284:22568–22579.

Iwasaki YW, Siomi MC, Siomi H. 2015. PIWI-interacting RNA: Its biogenesis and functions. Annu Rev Biochem 84:405–433.

Jawdekar GW, Hanzlowsky A, Hovde SL, Jelencic B, Feig M, Geiger JH, Henry RW. 2006. The unorthodox SNAP50 zinc finger domain contributes to cooperative promoter recognition by human SNAPC. J Biol Chem 281:31050–31060.

Kamath RS, Fraser AG, Dong Y, Poulin G, Durbin R, Gotta M, Kanapin A, Le Bot N, Moreno S, Sohrmann M et al. 2003. Systematic functional analysis of the *Caenorhabditis elegans* genome using RNAi. Nature 421:231–237.

Kasper DM, Wang G, Gardner KE, Johnstone TG, Reinke V. 2014. The *C. elegans* SNAPc component SNPC-4 coats piRNA domains and is globally required for piRNA abundance. Dev Cell 31:145–158.

Kirshner JA, Picard CL, Weiser NE, Mehta N, Feng S, Murphy VN, Vakhnovetsky A, Alessi AF, Xiao C, Inoki K, et al. 2025. Regulation of MORC-1 is key to the CSR-1-mediated germline gene licensing mechanism in *C. elegans*. Sci Adv 11:eado4170.

Kiuchi T, Koga H, Kawamoto M, Shoji K, Sakai H, Arai Y, Ishihara G, Kawaoka S, Sugano S, Shimada T et al. 2014. A single female-specific piRNA is the primary determiner of sex in the silkworm. Nature 509:633–636.

Kumar K, Trzybulska D, Tsatsanis C, Giwercman A, Almstrup K. 2019. Identification of circulating small non-coding RNAs in relation to male subfertility and reproductive hormones. Mol Cell Endocrinol 492:110443.

Lai H-T, Kang YS, Stumph WE. 2008. Subunit stoichiometry of the *Drosophila melanogaster* small nuclear RNA activating protein complex (SNAPc). FEBS Lett 582:3734–3738.

Langmead B, Trapnell C, Pop M, Salzberg SL. 2009. Ultrafast and memory-efficient alignment of short DNA sequences to the human genome. Genome Biol 10:R25.

Letunic I, Bork P. 2021. Interactive Tree Of Life (iTOL) v5: an online tool for phylogenetic tree display and annotation. Nucleic Acids Res 49:W293–W296.

Li C, Harding GA, Parise J, McNamara-Schroeder KJ, Stumph WE. 2004. Architectural arrangement of cloned proximal sequence element-binding protein subunits on *Drosophila* U1 and U6 snRNA gene promoters. Mol Cell Biol 24:1897–1906.

Li H, Handsaker B, Wysoker A, Fennell T, Ruan J, Homer N, Marth G, Abecasis G, Durbin R, 1000 Genome Project Data Processing Subgroup. 2009. The sequence alignment/map format and SAMtools. Bioinformatics 25:2078–2079.

Lin H, Spradling AC. 1997. A novel group of pumilio mutations affects the asymmetric division of germline stem cells in the *Drosophila* ovary. Development 124:2463–2476.

Luteijn MJ, Ketting RF. 2013. PIWI-interacting RNAs: from generation to transgenerational epigenetics. Nat Rev Genet 14:523–534.

Ma B, Hernandez N. 2001. A map of protein-protein contacts within the small nuclear RNA-activating protein complex SNAPc*. J Biol Chem 276:5027–5035.

Ma B, Hernandez N. 2002. Redundant cooperative interactions for assembly of a human U6 transcription initiation complex. Mol Cell Biol 22:8067–8078.

Mittal V, Ma B, Hernandez N. 1999. SNAPc: a core promoter factor with a built-in DNA-binding damper that is deactivated by the Oct-1 POU domain. Genes Dev 13:1807–1821.

Ozata DM, Gainetdinov I, Zoch A, O’Carroll D, Zamore PD. 2019. PIWI-interacting RNAs: small RNAs with big functions. Nat Rev Genet 20:89–108.

Paix A, Folkmann A, Rasoloson D, Seydoux G. 2015. High efficiency, homology-directed genome editing in *Caenorhabditis elegans* using CRISPR-Cas9 ribonucleoprotein complexes. Genetics 201:47–54.

Paniagua N, Roberts CJ, Gonzalez LE, Monedero-Alonso D, Reinke V. 2024. The Upstream Sequence Transcription Complex dictates nucleosome positioning and promoter accessibility at piRNA genes in the *C. elegans* germ line. PLoS Genet 20:e1011345.

Phillips CM, Montgomery BE, Breen PC, Roovers EF, Rim Y-S, Ohsumi TK, Newman MA, van Wolfswinkel JC, Ketting RF, Ruvkun G et al. 2014. MUT-14 and SMUT-1 DEAD box RNA helicases have overlapping roles in germline RNAi and endogenous siRNA formation. Curr Biol 24:839–844.

Ramírez F, Ryan DP, Grüning B, Bhardwaj V, Kilpert F, Richter AS, Heyne S, Dündar F, Manke T. 2016. deepTools2: a next generation web server for deep-sequencing data analysis. Nucleic Acids Res 44:W160–5.

Reinke V, Smith HE, Nance J, Wang J, Van Doren C, Begley R, Jones SJ, Davis EB, Scherer S, Ward S et al. 2000. A global profile of germline gene expression in *C. elegans*. Mol Cell 6:605– 616.

Robinson MD, McCarthy DJ, Smyth GK. 2010. edgeR: a Bioconductor package for differential expression analysis of digital gene expression data. Bioinformatics 26:139–140.

Ruby JG, Jan C, Player C, Axtell MJ, Lee W, Nusbaum C, Ge H, Bartel DP. 2006. Large-scale sequencing reveals 21U-RNAs and additional microRNAs and endogenous siRNAs in *C. elegans*. Cell 127:1193–1207.

Seroussi U, Lugowski A, Wadi L, Lao RX, Willis AR, Zhao W, Sundby AE, Charlesworth AG, Reinke AW, Claycomb JM. 2023. A comprehensive survey of *C. elegans* argonaute proteins reveals organism-wide gene regulatory networks and functions. Elife 12. doi:10.7554/eLife.83853

Shirayama M, Seth M, Lee H-C, Gu W, Ishidate T, Conte D Jr, Mello CC. 2012. piRNAs initiate an epigenetic memory of nonself RNA in the *C. elegans* germline. Cell 150:65–77.

Shi Z, Montgomery TA, Qi Y, Ruvkun G. 2013. High-throughput sequencing reveals extraordinary fluidity of miRNA, piRNA, and siRNA pathways in nematodes. Genome Res 23:497–508.

Simon M, Sarkies P, Ikegami K, Doebley A-L, Goldstein LD, Mitchell J, Sakaguchi A, Miska EA, Ahmed S. 2014. Reduced insulin/IGF-1 signaling restores germ cell immortality to *Caenorhabditis elegans* Piwi mutants. Cell Rep 7:762–773.

Spracklin G, Fields B, Wan G, Becker D, Wallig A, Shukla A, Kennedy S. 2017. The RNAi inheritance machinery of *Caenorhabditis elegans*. Genetics 206:1403–1416.

Sun J, Li X, Hou X, Cao S, Cao W, Zhang Y, Song J, Wang M, Wang H, Yan X et al. 2022. Structural basis of human SNAPc recognizing proximal sequence element of snRNA promoter. Nat Commun 13:6871.

Thomas J, Lea K, Zucker-Aprison E, Blumenthal T. 1990. The spliceosomal snRNAs of *Caenorhabditis elegans*. Nucleic Acids Res 18:2633–2642.

Vagin VV, Sigova A, Li C, Seitz H, Gvozdev V, Zamore PD. 2006. A distinct small RNA pathway silences selfish genetic elements in the germline. Science 313:320–324.

Wang G, Reinke V. 2008. A *C. elegans* Piwi, PRG-1, regulates 21U-RNAs during spermatogenesis. Curr Biol 18:861–867.

Wang X, Zeng C, Liao S, Zhu Z, Zhang J, Tu X, Yao X, Feng X, Guang S, Xu C. 2021. Molecular basis for PICS-mediated piRNA biogenesis and cell division. Nat Commun 12:5595.

Wang Y-H, Hertz HL, Pastore B, Tang W. 2025. An AT-hook transcription factor promotes transcription of histone, spliced-leader, and piRNA clusters. Nucleic Acids Res 53:gkaf079.

Weick E-M, Sarkies P, Silva N, Chen RA, Moss SMM, Cording AC, Ahringer J, Martinez-Perez E, Miska EA. 2014. PRDE-1 is a nuclear factor essential for the biogenesis of Ruby motif-dependent piRNAs in *C. elegans*. Genes Dev 28:783–796.

Weiser NE, Yang DX, Feng S, Kalinava N, Brown KC, Khanikar J, Freeberg MA, Snyder MJ, Csankovszki G, Chan RC et al. 2017. MORC-1 integrates nuclear RNAi and transgenerational chromatin architecture to promote germline immortality. Dev Cell 41:408–423.e7.

Weng C, Kosalka J, Berkyurek AC, Stempor P, Feng X, Mao H, Zeng C, Li W-J, Yan Y-H, Dong M-Q, Morero NR, Zuliani C, Barabas O, Ahringer J, Guang S, Miska EA. 2019. The USTC co-opts an ancient machinery to drive piRNA transcription in *C. elegans*. Genes Dev 33:90–102.

Williams Z, Morozov P, Mihailovic A, Lin C, Puvvula PK, Juranek S, Rosenwaks Z, Tuschl T. 2015. Discovery and characterization of piRNAs in the human fetal ovary. Cell Rep 13:854–863.

Wong MW, Henry RW, Ma B, Kobayashi R, Klages N, Matthias P, Strubin M, Hernandez N. 1998. The large subunit of basal transcription factor SNAPc is a Myb domain protein that interacts with Oct-1. Mol Cell Biol 18:368–377.

Yang Q, Hua J, Wang L, Xu B, Zhang H, Ye N, Zhang Z, Yu D, Cooke HJ, Zhang Y et al. 2013. MicroRNA and piRNA profiles in normal human testis detected by next generation sequencing. PLoS One 8:e66809.

Zanin E, Dumont J, Gassmann R, Cheeseman I, Maddox P, Bahmanyar S, Carvalho A, Niessen S, Yates JR 3rd, Oegema K et al.. 2011. Affinity purification of protein complexes in C. elegans. Methods Cell Biol 106:289–322.

Zeng C, Weng C, Wang X, Yan Y-H, Li W-J, Xu D, Hong M, Liao S, Dong M-Q, Feng X et al. 2019. Functional proteomics identifies a PICS complex required for piRNA maturation and chromosome segregation. Cell Rep 27:3561–3572.e3.

Zhang Y, Liu T, Meyer CA, Eeckhoute J, Johnson DS, Bernstein BE, Nusbaum C, Myers RM, Brown M, Li W et al.. 2008. Model-based analysis of ChIP-Seq (MACS). Genome Biol 9:R137.

Zhou X, Zuo Z, Zhou F, Zhao W, Sakaguchi Y, Suzuki Takeo, Suzuki Tsutomu, Cheng H, Zhou R. 2010. Profiling sex-specific piRNAs in zebrafish. Genetics 186:1175–1185.

Zhu C, Si X, Hou X, Xu P, Gao J, Tang Y, Weng C, Xu M, Yan Q, Jin Q, Cheng J, Ruan K, Zhou Y, Shan G, Xu D, Chen X, Xiang S, Huang X, Feng X, Guang S. 2025. piRNA gene density and SUMOylation organize piRNA transcriptional condensate formation. Nat Struct MolBiol 32:1503– 1516.

